# *De novo* and scaffold-based design of GDF15 binders for cancer cachexia diagnostics and therapeutics

**DOI:** 10.1101/2025.09.03.673894

**Authors:** Jinsook Ahn, Ryeongeun Cho, Sohyun Kim, Dongsun Lee, Ho Min Kim

## Abstract

Growth differentiation factor 15 (GDF15), a stress-responsive cytokine of the TGF-β superfamily, is elevated in cancer cachexia, chemotherapy-induced nausea, and hyperemesis gravidarum, making it both a biomarker and a therapeutic target. Here, we developed high-affinity GDF15 binders using an AI-driven protein design framework. To achieve this, we systematically explored three complementary scaffold-generation strategies: scaffold grafting (SG), diffusion-based *de novo* design, and scaffold-search and grafting (SSG), identifying distinct advantages - SG rapidly optimized receptor-derived motifs to subnanomolar affinity; *de novo* diffusion produced topologically novel binders; and SSG enabled access to GDF15’s concave site B by repurposing evolutionary structural analogs from natural complexes. The designed GDF15 binders were translated into two functional modalities. First, a one-step, wash-free luminescent biosensor was created by coupling a *de novo* binder to split-luciferase fragments, enabling the rapid and sensitive quantification of GDF15. Second, the highest-affinity binder was engineered as an Fc-fusion decoy receptor, thereby effectively neutralizing GDF15 signaling in cell-based assays (IC_50_ = 7.2 nM), demonstrating comparable *in vitro* potency to ponsegromab, a monoclonal antibody currently undergoing Phase II clinical trials. Together, this work establishes a versatile AI-driven binder design pipeline with broad potential for next-generation diagnostics and therapeutics in cancer cachexia and other GDF15-mediated diseases.

## 1. Introduction

Growth differentiation factor-15 (GDF15) is a stress-responsive hormone that belongs to the transforming growth factor-β (TGF-β) superfamily^1–3^. GDF15 is initially produced as a precursor (pre-pro-GDF15) and undergoes proteolytic processing to form a biologically active 25-kDa homodimer^4,5^. Under normal physiological conditions, circulating levels of GDF15 are low (0.1–1.2 ng/mL), and it plays a crucial role in the regulation of energy homeostasis, appetite, and body weight^1,6–8^. GDF15 exerts its biological functions by specifically binding to the glial cell-derived neurotrophic factor family receptor alpha-like (GFRAL), which is almost exclusively expressed in neurons of the area postrema (AP) and nucleus tractus solitarius (NTS) in the hindbrain^9,10^. Upon GDF15 binding, GFRAL recruits the receptor tyrosine kinase RET, thereby activating downstream pathways that regulate metabolism and energy expenditure (Fig. 1a)^11–13^.

**Figure 1.**
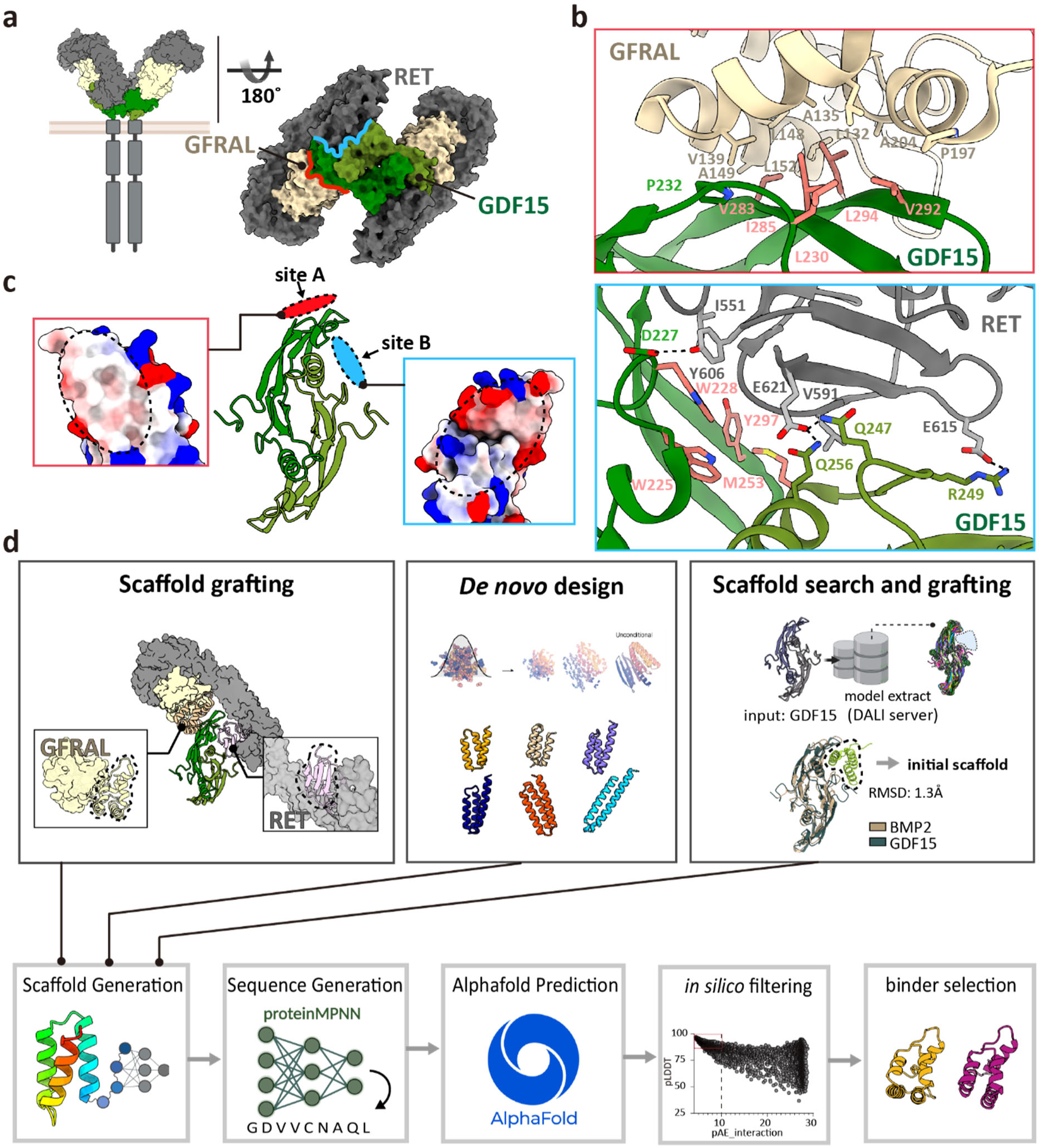
Structure of GDF15/GFRAL/RET complex and design strategy for GDF15 antagonistic binders. **a.** Cryo-EM structure of the extracellular GDF15–GFRAL–RET complex (PDB: 6Q2J) showing a 2:2:2 stoichiometry, wherein the dimeric GDF15 bridges two GFRAL co-receptors and two RET receptors. **b.** Binding interfaces of GDF15 with GFRAL (top) and RET (bottom). Key interacting residues are shown as sticks. Hydrophobic hotspot residues used for binder design are highlighted in pink. **c.** Target sites for GDF15 binder design. The convex surface engaging GFRAL (site A, red) and the concave surface contacting RET (site B, light blue) are highlighted. Insets show electrostatic surface potentials of each site (white, hydrophobic; blue, positive charge; red, negative charge). **d.** Workflow of binder design using three scaffold generation strategies: Scaffold Grafting (SG), Diffusion-based *de novo* Design, and Scaffold-Search and Grafting (SSG).

Pathologically, circulating GDF15 levels can increase dramatically, often reaching 10–100 ng/ml or higher, in response to acute injury, chronic inflammation, cardiovascular disease, pregnancy complications, and diverse malignancies^14^. Elevated GDF15 exerts anorexigenic effects, leading to reduced food intake, muscle wasting, and severe weight loss, while also stimulating the peripheral sympathetic nervous system^15,16^. In particular, high GDF15 levels are strongly correlated with cancer cachexia, a multifactorial syndrome characterized by anorexia, skeletal muscle atrophy, and metabolic dysfunction ^2,17,18^. Cachexia affects more than 50% of patients with advanced cancer, profoundly diminishing quality of life, reducing responsiveness to chemotherapy, and contributing to cancer-related mortality^19^. Notably, cisplatin-based chemotherapy induces acute GDF15 elevation, which has been directly linked to chemotherapy-induced nausea and vomiting^16,17^. Despite its clinical impact, cachexia remains largely untreatable, underscoring the urgent need for early diagnostic tools and targeted therapies.

Given the association between GDF15 and cachexia, recent diagnostic and therapeutic efforts have predominantly focused on antibody-based strategies targeting GDF15^18,20,21^. Pfizer’s ponsegromab (PF-06946860), a monoclonal antibody against GDF15, has demonstrated clinical efficacy. In a recent Phase 2 randomized, double-blind clinical trial in patients with cancer cachexia and elevated GDF15 levels—including those with non-small cell lung cancer, pancreatic cancer, and colorectal cancer— ponsegromab treatment led to significant weight gain, improved appetite, increased physical activity, and reduced cachexia symptoms compared to placebo, while maintaining a favorable safety profile^20,22^. Similarly, Roche’s Elecsys GDF15 assay, approved by the FDA as a companion diagnostic, enables precise monitoring of circulating GDF15 levels to support disease diagnosis and treatment decision-making.

In this study, we leverage recent breakthroughs in AI-based protein design to develop synthetic minibinders that selectively target the receptor-binding surface of GDF15 with high affinity. The initial selection of the minibinder scaffold is a critical determinant of design success; however, conventional diffusion-based approaches require extensive candidate screening and often fail to yield effective binders for targets with highly polar surfaces. To overcome these challenges, we implemented and compared three complementary scaffold design strategies for the development of a GDF15 binder: (1) scaffold grafting (SG), (2) diffusion-based *de novo* design, and (3) scaffold-search and grafting (SSG). Using these designed binders targeting GDF15, we developed highly sensitive biosensors capable of rapid one-step detection of GDF15 at nanomolar concentrations, as well as effective therapeutic blockers that neutralize GDF15 activity, offering novel strategies for the effective diagnosis and treatment of cancer cachexia.

## 2. Results

### 2.1. Structural analysis of GDF15/GFRAL/RET complex and rational design strategy for GDF15 antagonistic binders

The cryo-electron microscopy (Cryo-EM) structure of the extracellular GDF15/GFRAL/RET complex reveals a 2:2:2 stoichiometry, wherein the dimeric GDF15 is centrally located, symmetrically bridging two GFRAL co-receptors and two RET receptors^23^. This characteristic batwing-shaped architecture brings the membrane-proximal cysteine-rich domains (CRDs) of RET into close proximity, thereby facilitating intracellular kinase dimerization for signal activation (Fig. 1a). Each GDF15 protomer interacts with the D2 domain of GFRAL (GFRAL D2) via the convex surface of its finger loops (site A) and with the cysteine-rich domains (CRD) of RET through the opposite concave surface (site B) (Fig. 1a–c), effectively wedging the GDF15 dimer between GFRAL and RET. This dual engagement explains the cooperative requirement of both receptors for GDF15-mediated signaling.

Based on these structural insights into the GDF15/GFRAL/RET complex, we designed antagonistic binders to block either site A or site B. Site A forms a smooth, hydrophobic-rich convex surface, whereas site B exhibits a concave topology with fewer hydrophobic patches (Fig. 1b–c). Key hydrophobic residues at site A (L230, V283, I285, V292, L294) and site B (W225, W228, M253, Y297) were designated as binding hotspots for binder design (Fig. 1b). To generate candidate scaffolds, we applied three complementary strategies: (1) scaffold grafting (SG), (2) diffusion-based *de novo* design using RFdiffusion, and (3) scaffold-search and grafting (SSG) (Fig. 1d). For each backbone, amino acid sequences were generated using ProteinMPNN^24^, and their complex structures with the GDF15 dimer were predicted with AF2 initial guess^25^ and AF3^26^. Designed binders were subsequently evaluated *in silico* based on predicted aligned error (pAE), predicted local-distance difference test (pLDDT) scores, and predicted changes in binding free energy (ΔΔG). These filtering metrics facilitated the identification of the most promising candidates, which exhibited high structural integrity and advantageous predicted binding characteristics (Fig. 1d).

### 2.2 Design of GDF15 site A binders via scaffold grafting-based backbone generation

To generate antagonistic binders targeting site A of GDF15, we initially employed a scaffold grafting (SG) approach utilizing a structural segment of the co-receptor GFRAL (Fig. 2a). Specifically, the triangular-shaped D2 domain of GFRAL (GFRAL D2, residues 129–211), which directly engages GDF15 site A, consists of five α-helices stabilized by five intramolecular disulfide bonds (Fig. 2b). This domain was extracted and employed as the initial scaffold for binder design.

**Figure 2.**
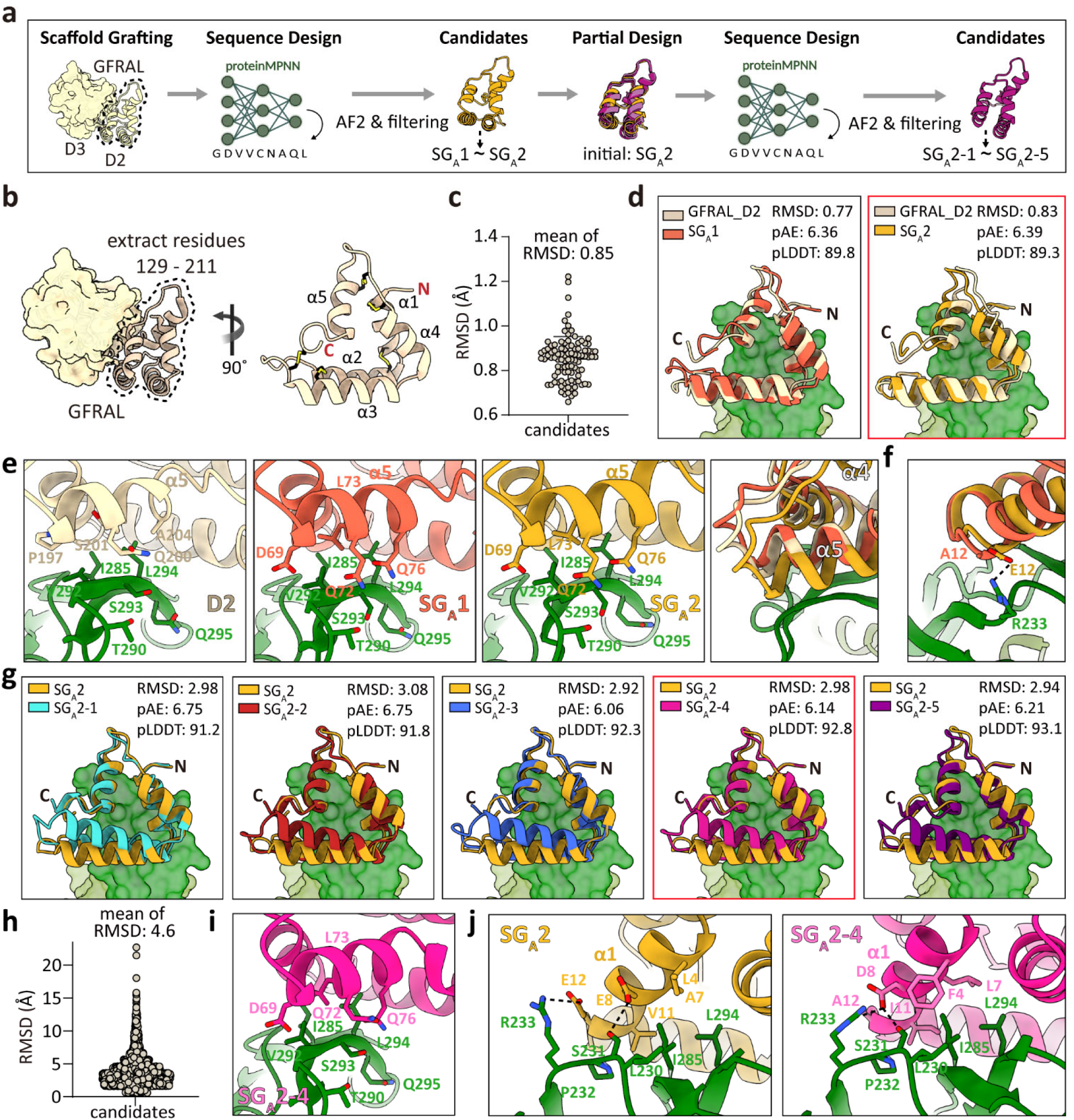
Design and optimization of GDF15 site A binders via scaffold grafting (SG). a. Workflow of binder design via scaffold grafting. The GFRAL D2 domain was extracted as the initial scaffold to generate SG_A_1 and SG_A_2. SG_A_2 was further optimized by scaffold-guided partial diffusion, resulting in five variants (SG_A_2-1 to SG_A_2-5). b. Extracted GFRAL D2 scaffold (residues 129∼211) from the GFRAL extracellular domain. The N- and C-termini, helices, and disulfide bonds are indicated. c. Backbone RMSD distribution of 100 ProteinMPNN-designed variants relative to the parental GFRAL D2 scaffold (mean RMSD = 0.85 Å). d. Structural alignment of the initial scaffold GFRAL D2 with SG_A_1 (left) and SG_A_2 (right). RMSD with AF3-predicted structure and AF3 scores (pAE_interaction and pLDDT) for the binder/GDF15 complex are indicated, with the N- and C-termini labeled. e. Structural comparison of GFRAL-D2, SG_A_1, and SG_A_2 at the binding interface. Residues that enhance binding, commonly observed in both SG_A_1 and SG_A_2, are indicated. The rightmost panel shows the superposition of three scaffolds, highlighting α5 displacement. f. Superposition of SG_A_1 and SG_A_2, showing a unique electrostatic interaction in SG_A_2. g. Structural alignment of SG_A_2 with its partial diffusion-derived variants. RMSD with AF2-predicted structure and AF2 scores (pAE_interaction and pLDDT) for the binder/GDF15 complex are indicated, with the N- and C-termini labeled. h. Backbone RMSD distribution of partial diffusion–derived variants relative to the SG_A_2 scaffold (mean RMSD = 4.6 Å). i. Structural model of SG_A_2-4 at the binding interface. Conserved binding residues in α5 (D69, Q72, L73, Q76), retained in SG_A_2-4 variants compared to SG_A_2, are highlighted. j. Binding interface comparison of SG_A_2 and SG_A_2-4, highlighting the shifted α1 position and distinct interacting residues on this helix.

Using this scaffold, 100 sequence variants were generated with ProteinMPNN to stabilize the backbone and optimize the site A binding interface. Predicted structural models showed that most variants closely resembled the initial scaffold, maintaining a mean backbone RMSD of ∼0.85 Å and preserving the relative orientation of the N- and C-termini (Fig. 2c-d). From this pool (28%, 28 out of 100 designs, pLDDT > 85, pAE_interaction < 10, ΔΔG < –30), two leading candidates were selected based on the metrics of pAE_interaction, pLDDT, and ΔΔG for subsequent experimental evaluation (Fig. 2d and S1a).

Interestingly, both binders carried cysteine substitutions (C10L/C16A in SG_A_1, C10I in SG_A_2), which abolished the α1–α2 disulfide bond present in native GFRAL D2 (Fig. S1d). Nevertheless, AF3 predictions demonstrated that both variants maintained the triangular fold, although SG_A_2 exhibited slight deviations from SG_A_1 and GFRAL D2 (RMSD 0.77 Å for SG_A_1 and 0.83 Å for SG_A_2 relative to GFRAL D2) (Fig. 2d). In the case of SG_A_2, new salt bridges formed between residues E47–K59 (α5-adjacent loop) and K75–D81 (α3 to α4), resulting in the displacement of helices α4 and α5 (Fig. S1d). This structural rearrangement introduced subtle distortions to the triangular geometry of SG_A_2 compared to SG_A_1 and GFRAL D2, while still preserving the overall integrity of the scaffold (Fig. 2d).

When expressed in *Escherichia coli* (*E. coli*) and purified using Ni–NTA affinity and size-exclusion chromatography (SEC), both SG_A_1 and SG_A_2 exhibited high yield, solubility, and homogeneity (Fig. S1b-c). Binding analysis indicated that, compared to the initial scaffold (K_D_ = 90 nM), SG_A_1 and SG_A_2 achieved enhanced affinities with K_D_ values of 50 nM and 5 nM, respectively (Table 1 and Fig. S1e). Based on AF3 predictions, we speculate that this affinity enhancement resulted from hydrophobic core interactions between residue L73 (S201 in GFRAL D2) and GDF15 site A residues (I285, V292, and L294), together with electrostatic interactions mediated by D69, Q72, and Q76 (P197, Q200, and A204 in GFRAL D2) (Fig. 2e). In SG_A_2, the displacement of α5 brought this helix closer to the β-strand of GDF15 site A, thereby strengthening SG_A_2 Q76-GDF15 S293 main chain, SG_A_2 D69-GDF15 G291 main chain, and SG_A_2 Q72-GDF15 T290 interactions. Additionally, a new electrostatic contact between SG_A_2 E12 and GDF15 R233 further stabilized the SG_A_2-GDF15 interface (Fig. 2f). Although the atomic-resolution structure has yet to be determined, these interactions likely account for the ∼10-fold higher affinity of SG_A_2 compared to SG_A_1.

**Table 1.** Binding kinetics summary for GFRAL D2 and designed GDF15 binders. Data from SPR analysis (Fig. S1e, S1i, S2c, and S3e) include association rates (kon1 and kon2), dissociation rates (koff1 and koff2), and equilibrium dissociation constant (K_D_). Binders with insufficient protein yield are marked as ‘–’, and binders with no detectable binding are marked as ‘ND’.

|  | Kon1 (1/Ms) | Koff1 (1/s) | Kon2 (1/s) | Koff2 (1/s) | K<sub>D</sub> (M) |
| --- | --- | --- | --- | --- | --- |
| **GFRAL D2** | 1.70x10<sup>5</sup> | 3.43x10<sup>-2</sup> | 3.10x10<sup>-3</sup> | 2.50x10<sup>-3</sup> | 9.04x10<sup>-8</sup> |
| **SG<sub>A</sub> 1** | 1.02x10<sup>6</sup> | 1.50x10<sup>-1</sup> | 3.46x10<sup>-3</sup> | 1.92x10<sup>-3</sup> | 5.24x10<sup>-8</sup> |
| **SG<sub>A</sub> 2** | 1.11x10<sup>6</sup> | 1.01x10<sup>-2</sup> | 3.99x10<sup>-3</sup> | 6.81x10<sup>-3</sup> | 5.81x10<sup>-9</sup> |
| **SG<sub>A</sub> 2-1** | 1.11x10<sup>6</sup> | 0.42x10<sup>-1</sup> | 0.18x10<sup>-2</sup> | 0.27x10<sup>-2</sup> | 2.32x10<sup>-8</sup> |
| **SG<sub>A</sub> 2-2** | - | - | - | - | - |
| **SG<sub>A</sub> 2-3** | 1.27x10<sup>6</sup> | 0.66x10<sup>-2</sup> | 0.17x10<sup>-1</sup> | 0.18x10<sup>-2</sup> | 4.99x10<sup>-10</sup> |
| **SG<sub>A</sub> 2-4** | 8.89x10<sup>6</sup> | 0.65x10<sup>-2</sup> | 0.35x10<sup>-2</sup> | 0.29x10<sup>-2</sup> | 3.30x10<sup>-10</sup> |
| **SG<sub>A</sub> 2-5** | 9.15x10<sup>3</sup> | 0.26x10<sup>-1</sup> | 0.26x10<sup>-1</sup> | 0.49x10<sup>-2</sup> | 4.66x10<sup>-7</sup> |
| **DE<sub>A</sub> 1** | - | - | - | - | - |
| **DE<sub>A</sub> 2** | - | - | - | - | - |
| **DE<sub>A</sub> 3** | 1.92x10<sup>6</sup> | 2.36x10<sup>-1</sup> | 1.13x10<sup>-1</sup> | 7.43x10<sup>-6</sup> | 7.59x10<sup>-9</sup> |
| **DE<sub>A</sub> 4** | 3.69x10<sup>6</sup> | 1.69x10<sup>-1</sup> | 1.35x10<sup>-4</sup> | 8.78x10<sup>-4</sup> | 3.97x10<sup>-8</sup> |
| **DE<sub>A</sub> 5** | 1.34x10<sup>5</sup> | 6.88x10<sup>-2</sup> | 5.64x10<sup>-5</sup> | 7.77x10<sup>-5</sup> | 2.97x10<sup>-7</sup> |
| **DE<sub>A</sub> 3-1** | 6.91x10<sup>5</sup> | 2.49x10<sup>-1</sup> | 5.97x10<sup>-7</sup> | 4.90x10<sup>-3</sup> | 3.61x10<sup>-7</sup> |
| **DE<sub>A</sub> 3-2** | 1.01x10<sup>6</sup> | 6.28x10<sup>-1</sup> | 1.36x10<sup>-6</sup> | 1.25x10<sup>-1</sup> | 6.23x10<sup>-7</sup> |
| **DE<sub>A</sub> 3-3** | - | - | - | - | - |
| **DE<sub>A</sub> 3-4** | - | - | - | - | - |
| **DE<sub>A</sub> 3-5** | 8.03x10<sup>6</sup> | 6.48x10<sup>-2</sup> | 1.47x10<sup>-3</sup> | 9.27x10<sup>-3</sup> | 6.96x10<sup>-9</sup> |
| **DE<sub>A</sub> 3-6** | - | - | - | - | - |
| **DE<sub>A</sub> 3-7** | - | - | - | - | - |
| **SSG<sub>B</sub>1** | - | - | - | - | - |
| **SSG<sub>B</sub>2** | 1.55x10<sup>5</sup> | 7.16x10<sup>-2</sup> | 1.51x10<sup>-2</sup> | 5.36x10<sup>-3</sup> | 1.21x10<sup>-7</sup> |
| **SSG<sub>B</sub>3** | ND | ND | ND | ND | ND |
| **SSG<sub>B</sub>4** | - | - | - | - | - |
| **SSG<sub>B</sub>5** | ND | ND | ND | ND | ND |

To further improve the binding affinity of SG_A_2, we applied partial diffusion in RFdiffusion with site A hotspots (L230, V283, I285, V292, and L294) specified as constraints and conducted cysteine-free sequence design (Fig. 2a). From a filtered pool, five leading variants (SG_A_2-1 to SG_A_2-5) were selected through *in silico* filtering (see Methods) (Fig. 2a and S1f). These variants exhibited 34–42% sequence identity to the parental scaffold SG_A_2, and a significant structural rearrangement of α3, which shifted by ∼9 Å (Fig. 2g). By eliminating disulfide bonds during sequence design, the scaffold’s rigid triangular form gained increased flexibility, allowing it to adopt shapes more complementary to GDF15 site A and thereby increasing backbone diversity (Fig. 2h).

Notably, among these candidates, SG_A_2-3 and SG_A_2-4 exhibited approximately 10-to 17-fold higher binding affinity than parental SG_A_2 (SG_A_2 K_D_ = 5 nM; SG_A_2-3 K_D_ = 500 pM; SG_A_2-4 K_D_ = 300 pM) (Table 1 and Fig. S1g-i), whereas SG_A_2-2 was expressed but aggregated, precluding affinity measurement. Although the sequence identity between SG_A_2 and SG_A_2-4 is only 31.3%, the key interacting residues in α5 (D69, Q72, L73, and Q76) were conserved (Fig. 2i). In contrast, several GDF15-interacting residues in α1 were altered, and the slight difference in α1 between SG_A_2 and SG_A_2-4 may account for the higher binding affinity of SG_A_2-4 (Fig. 2j).

Collectively, scaffold grafting followed by a single round of partial diffusion and sequence redesign yielded a high-affinity GDF15 binder, exhibiting a 300-fold enhancement in binding affinity compared to the original GFRAL D2 scaffold.

### 2.3 Design of GDF15 site A binders via diffusion-based *de novo* backbone generation

To further investigate structural diversity beyond scaffold grafting, we employed a diffusion-based *de novo* backbone generation strategy using RFdiffusion (Fig. 3a). Protein backbones comprising 50 to 90 amino acids were generated with site A hotspots (L230, V283, I285, V292, and L294) specified as constraints, resulting in the production of 1,728 scaffolds. All outputs generated by RFdiffusion formed helical bundles consisting of one to five helices, predominantly adopting two- or three-helix bundles (Fig. 3b). For each backbone, three sequence variants were designed, resulting in 5,178 unique sequences. These were evaluated by AF2 predictions and computational filtering (Fig. S2a). From this pool (10.4%, 539 out of 5,184 designs, pLDDT > 85, pAE_interaction < 10, ΔΔG < –30), five top-ranked candidates (DE_A_1– DE_A_5) were selected for experimental validation (Fig. 3c). Structural modeling showed that all five candidates formed three-helix bundles covering the entire surface of GDF15 site A, with their N- and C-termini oriented in opposite directions (Fig. 3c).

**Figure 3:**
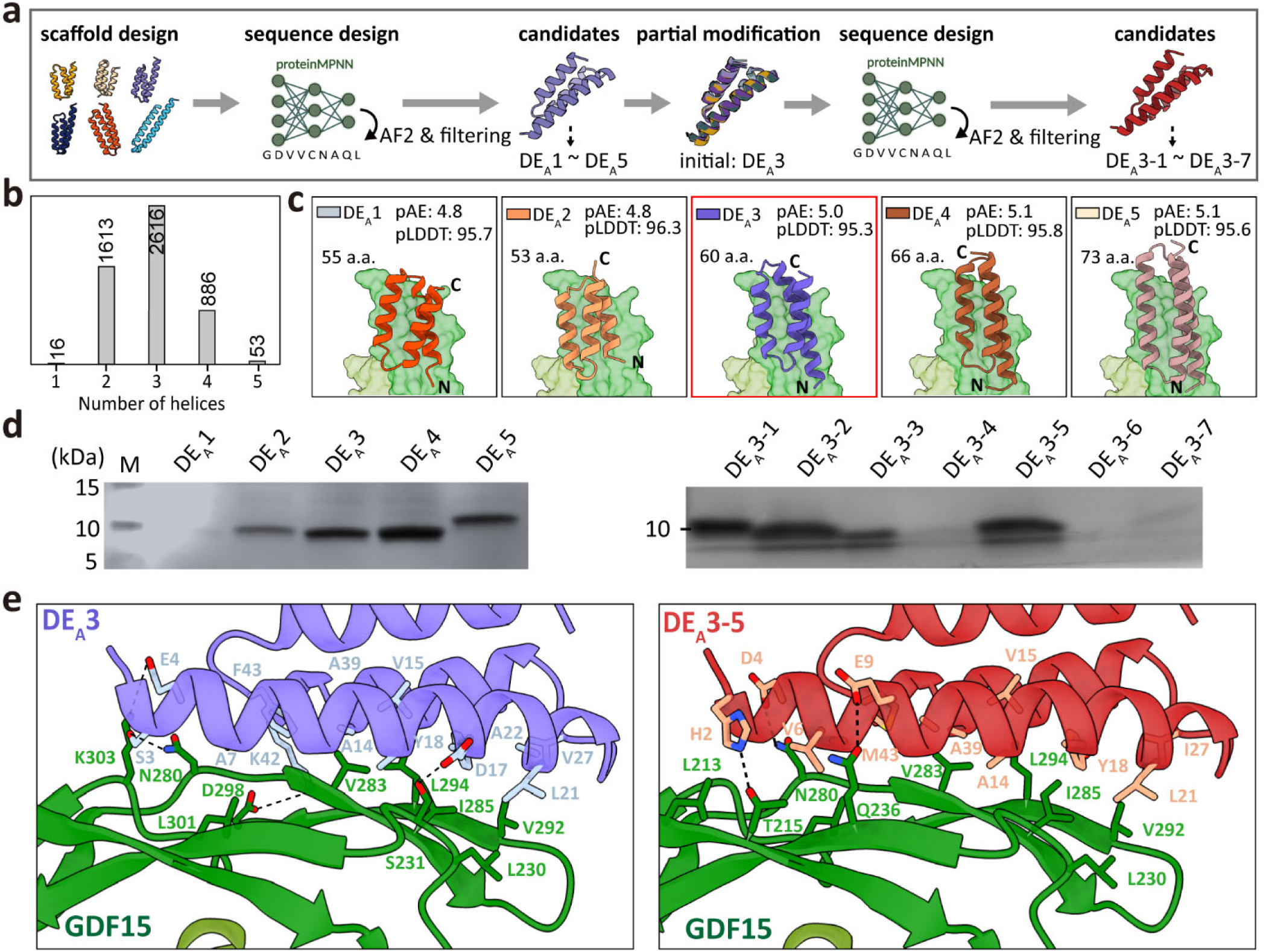
Design and optimization of GDF15 site A binders via diffusion-based *de novo* design. a. Workflow of *de novo* binder design using RF diffusion. A total of 1,728 initial scaffolds (50∼90 a.a) were generated, sequence-designed (three sequences per each backbone), and computationally filtered using *in silico* evaluation metrics. The top five binders (DE_A_1–DE_A_5) were structurally analyzed and experimentally validated by expression, purification, and binding analysis. The best-performing DE_A_3 was further optimized by scaffold-guided partial diffusion, resulting in seven variants (DE_A_3-1 to DE_A_3-7). b. Distribution of helix counts in RFdiffusion-generated scaffolds to analyze structural diversity. c. AF2-predicted structural models of the five selected *de novo* binder candidates in complex with the GDF15 dimer. Binder lengths, pAE_interaction, and pLDDT values are indicated, with the N- and C-termini labeled. d. SDS-PAGE analysis of binders (DE_A_1–5, left; DE_A_3-1 to DE_A_3-7, right) after *E. coli* expression and affinity purification. e. Binding interface comparison of DE_A_3 (left) and DE_A_3-5 (right) with GDF15. Key interacting residues are shown as sticks and labeled.

Expression screening in *E. coli* showed that DE_A_3, DE_A_4, and DE_A_5 were soluble and purified with high homogeneity, while DE_A_1 failed to express and DE_A_2 aggregated during purification (Fig. 3d, left and S2b). Despite favorable *in silico* scores (low pAE_interaction and high pLDDT scores), variations in expression and solubility were pronounced. Additionally, binding affinities ranged from low nanomolar to hundreds of nanomolar (DE_A_3, K_D_ = 7.6 nM; DE_A_4, K_D_ = 39 nM; and DE_A_5, K_D_ = 297 nM) (Table 1 and Fig. S2c). These findings underscore the limitations of predictive metrics in comprehensively capturing the physiochemical properties of designed binders. DE_A_3 achieved the strongest binding through extensive hydrophobic contacts (A7, A14, V15, Y18, L21, A22, V27, A39, and F43 of DE_A_3 with L230, V283, I285, V292, and L294 of GDF15 site A). The binding was further stabilized by surrounding electrostatic interactions (DE_A_3 S3-GDF15 N280, DE_A_3 E4-GDF15 K303, DE_A_3 D17-GDF15 S231, and DE_A_3 K42-GDF15 D298). (Fig. 3e, left).

Similar to the affinity maturation of the SG_A_2 binder described above, we employed DE_A_3-guided partial diffusion, followed by sequence design, structural prediction, and computational filtering for the affinity maturation of DE_A_3 binder (Fig. 3a). Out of a total of 6,000 candidates, the top 100 candidates (pAE_interaction, 4.3–5.0; pLDDT, 94–97) were selected and further screened experimentally using yeast surface display (Fig. S3a). Fluorescence-activated cell sorting (FACS) with dual labeling (anti-Myc PE-Cy7 for binder expression and PE-conjugated Strep-Tactin for GDF15 binding) facilitated the recovery of the top 25% double-positive cells, and sequencing of these cells identified seven unique candidates (DE_A_3-1 to DE_A_3-7). The sequence identities of DE_A_3-derived variants (DE_A_3-1 to DE_A_3-7) ranged from 46% to 58%, although their structures remained highly conserved (RMSDs 0.6–2.0 Å) (Fig. S3b-c).

Among the seven DE_A_3-derived candidates, only three (DE_A_3-1, DE_A_3-2, and DE_A_3-5) were successfully expressed and purified in *E. coli* (Fig. 3d, right, and S3d). Notably, unlike scaffold grafting– based backbone generation followed by partial diffusion, where SG_A_2-3 and SG_A_2-4 enhanced affinity by 10 to 17-fold, DE_A_3-derived design variants did not exceed the performance of the parental DE_A_3 (DE_A_3, K_D_ = 7.5 nM; DE_A_3-1, K_D_ = 361 nM; DE_A_3-2, K_D_ = 623 nM; and DE_A_3-5, K_D_ = 6.9 nM) (Table 1 and S3e).

Structural analysis confirmed that DE_A_3-5 recapitulated the binding mode of DE_A_3, engaging the entire surface of site A on GDF15 through a hydrophobic core network further stabilized by neighboring electrostatic interactions (Fig. 3e, right). Although the affinities of DE_A_3 and DE_A_3-5 were ∼20-fold weaker than that of SG_A_2-4, their three-helix bundle structure with oppositely oriented termini provided a distinct advantage for biosensor development (see Section 2.5).

Taken together, diffusion-based *de novo* backbone generation successfully produced nanomolar-affinity binders and enabled the development of GDF15 site A binders with distinct topologies compared to those derived from scaffold grafting.

### 2.4 Design of GDF15 site B binders via scaffold-search and grafting for backbone generation

To design antagonistic binders targeting site B of GDF15, which has fewer exposed hydrophobic residues and more polar residues compared to site A (Fig. 1b–c), we initially assessed the RET receptor, which naturally interacts with this site. As described in section 2.2, a domain segment of RET (residues 586-622), comprising five β-strands stabilized by two disulfide bonds, was extracted as an initial scaffold. One hundred sequence variants were generated and subsequently evaluated using AlphaFold2 predictions (Fig. 4a). However, aside from the two disulfide-stabilized β-strands, most elements unfolded, indicating that this RET-derived scaffold lacked sufficient stability for further development (Fig. 4b).

**Figure 4:**
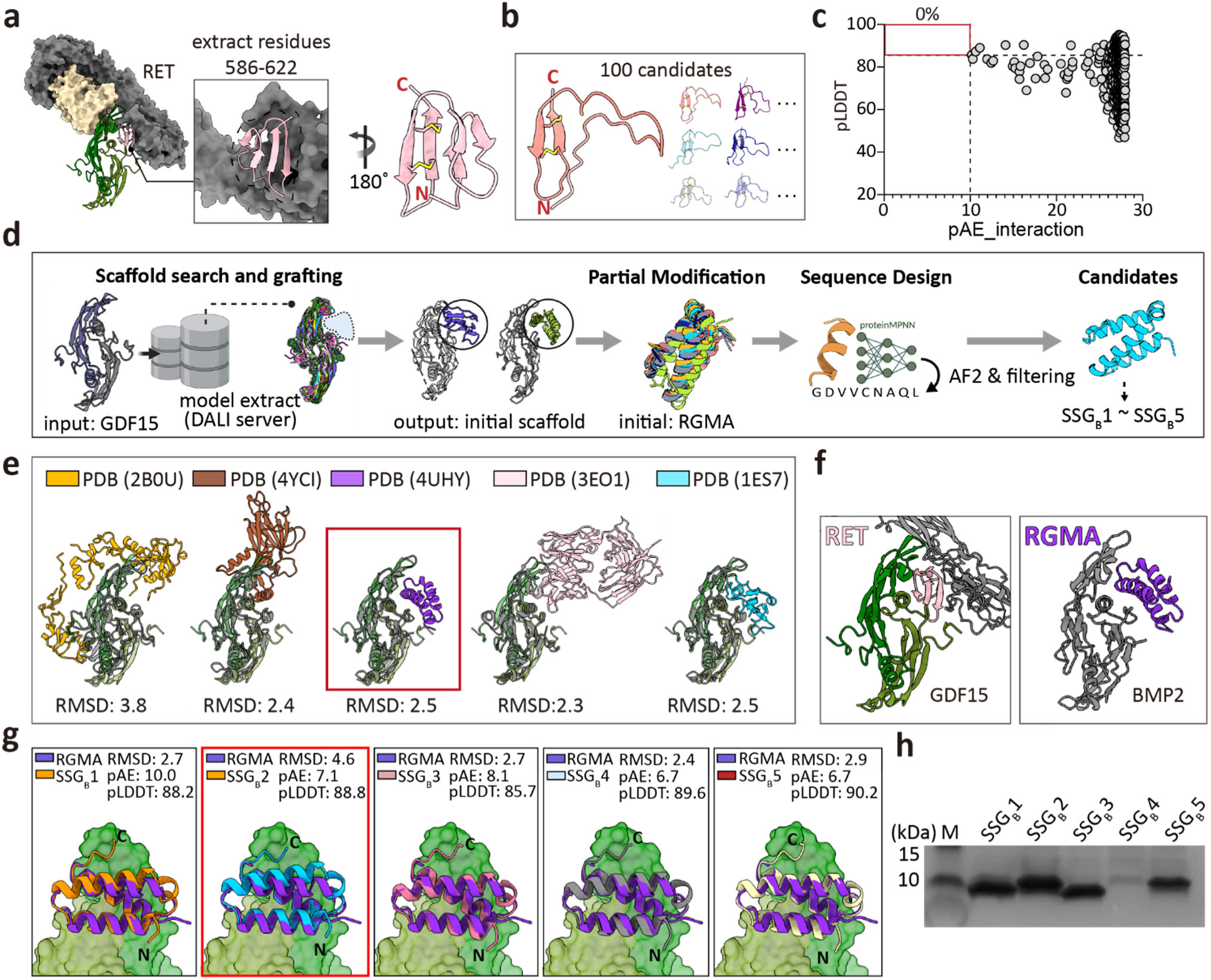
Design of GDF15 site B binders via scaffold-search and grafting (SSG). a. The RET domain segment (residues 586–622) was tested as an initial scaffold for site B binders design. b. AF2-predicted structures of 100 RET-derived variants by sequence design. Only disulfide-constrained β-strands remained folded. c. *In silico* filtering of 500 *de novo* backbones generated by RFdiffusion with site B hotspot constraints (W225, W228, M253, and Y297). No candidates satisfied filtering thresholds (pLDDT > 85 and pAE_interaction < 10; red box). d. Workflow of the scaffold search and grafting (SSG) strategy. The GDF15 structure was used as a query in the DALI server to search the Protein Data Bank (PDB) for natural scaffolds with similar topology and surface geometry. Candidate scaffolds were then subjected to scaffold-guided partial diffusion and sequence design. e. Representative scaffold candidates identified from the DALI server search: Follistatin/Activin A (PDB 2B0U), BMP9 pro-complex (mature domain + prodomain) (PDB 4YCI), BMP2/RGMA (PDB 4UHY), TGF-β3/GC-1008 antibody (PDB 3EO1), and BMP2/BMP2 receptor A (PDB 1ES7). RMSD relative to GDF15 is indicated. f. Structural comparison of GDF15/RET (green/pink) and BMP2/RGMA (olive/purple) complexes. GDF15 and BMP2 show overall similarity (RMSD = 2.5 Å), but interacting partners differ topologically. g. Structural alignment of RGMA with RGMA-derived binder variants (SSG_B_1 to SSG_B_5). RMSD with AF2-predicted structures and AF2 scores (pAE_interaction and pLDDT) for the binder/GDF15 complex are indicated. h. SDS-PAGE analysis of binders (SSG_B_1-SSG_B_5) after *E. coli* expression and affinity purification.

Next, we employed *de novo* scaffold generation using RFdiffusion and ProteinMPNN with site B hotspot constraints (W225, W228, M253, Y297; hydrophobic residues at site B), producing 100 backbones of 50–90 amino acids and five sequence variants per backbone (500 total designs). Nonetheless, none of these candidates fulfilled the *in silico* filtering criteria (pLDDT > 85, pAE_interaction < 10, ΔΔG < –30), thereby underscoring the challenges associated with designing stable site B binders through direct extraction or *de novo* diffusion-based methods (Fig. 4c).

To address these limitations, we developed a scaffold search and grafting (SSG) strategy aimed at identifying naturally stable scaffolds sharing comparable structural features (Fig. 4d). Using the GDF15 structure as a query, we searched the Protein Data Bank (PDB) using DALI to find proteins with similar topology and surface geometry, thereby leveraging evolutionary insights from natural complexes. From the search results obtained, structures in apo form (comprising solely GDF15-like folds) were excluded, while the top-ranking PDB entries in which GDF15-like folds participated in protein–protein interactions were retained. This search identified suitable scaffolds for accommodating GDF15 site B from the complex of Follistatin/Activin A [PDB 2B0U]^27^, BMP9 pro-complex (mature domain + prodomain) [PDB 4YCI]^28^, BMP2/RGMA [PDB 4UHY]^29^, TGF-β3/GC-1008 antibody [PDB 3EO1]^30^, and BMP2/BMP2 receptor A [PDB 1ES7]^31^ (Fig. 4e).

Among these, we selected RGMA in complex with BMP2 (PDB 4UHY) as the initial scaffold because it forms a compact and stable α-helical bundle, unlike the unstable RET-derived scaffold (Fig. 4b). Specifically, the GDF15 dimer was superimposed onto BMP2 in the BMP2/RGMA complex, and BMP2 was replaced with the aligned GDF15 dimer while preserving RGMA. Although BMP2 and GDF15 exhibit considerable structural similarity (RMSD = 2.5 Å), notable differences exist in their flexible loop regions (Fig. 4e-f). To account for these subtle variations, the resulting model was further optimized through scaffold-guided partial diffusion and sequence design, yielding a total of 3,000 designs (Fig. 4d). After filtering (0.3%, 10 out of 3000 designs, pLDDT > 85, pAE_interaction < 10, ΔΔG < –30), five top candidates (SSG_B_1–SSG_B_5) were selected for experimental testing (Fig. 4g and Fig. S4a). Although only about 0.3% of the designs passed the filters (10 of 3,000), the expression success rate was high at 80% (4 of 5), likely reflecting the stability of the natural scaffold (Fig. 4h and S4b). Among these, only SSG_B_2 bound GDF15 measurably, with a K_D_ of 121 nM (Table 1 and Fig. S4c). Structural prediction models suggested that two of the three helices in SSG_B_2 are positioned in direct contact with the concave binding surface at site B of GDF15. Although multiple electrostatic interactions are present across the binding interface, the limited availability of exposed hydrophobic residues at site B significantly restricts hydrophobic core formation between SSG_B_2 and GDF15 (Fig. S4d).

Collectively, the SSG strategy offers a promising framework for binder design, particularly when natural binders are unavailable or unstable, and the target surface lacks sufficient hydrophobic residues for effective *de novo* design. This methodology significantly broadens the array of tools available for developing binders against previously inaccessible targets through conventional approaches.

### 2.5. Development and optimization of a BAT biosensor using designed GDF15 binders for rapid detection

To enable rapid and cost-effective detection of GDF15, we adapted the binding-activated tandem split-enzyme (BAT) system, which was originally developed for single-component biosensing^32^. In this format, a high-affinity GDF15 binder is fused to the N- and C-terminal segments of NanoLuc luciferase (SmBiT and LgBiT). In the absence of GDF15, the luciferase fragments remain apart, preventing reconstitution. Upon GDF15 binding, steric rearrangement brings the fragments into proximity, restoring luciferase activity and producing a luminescent signal (Fig. 5a–b).

**Figure 5.**
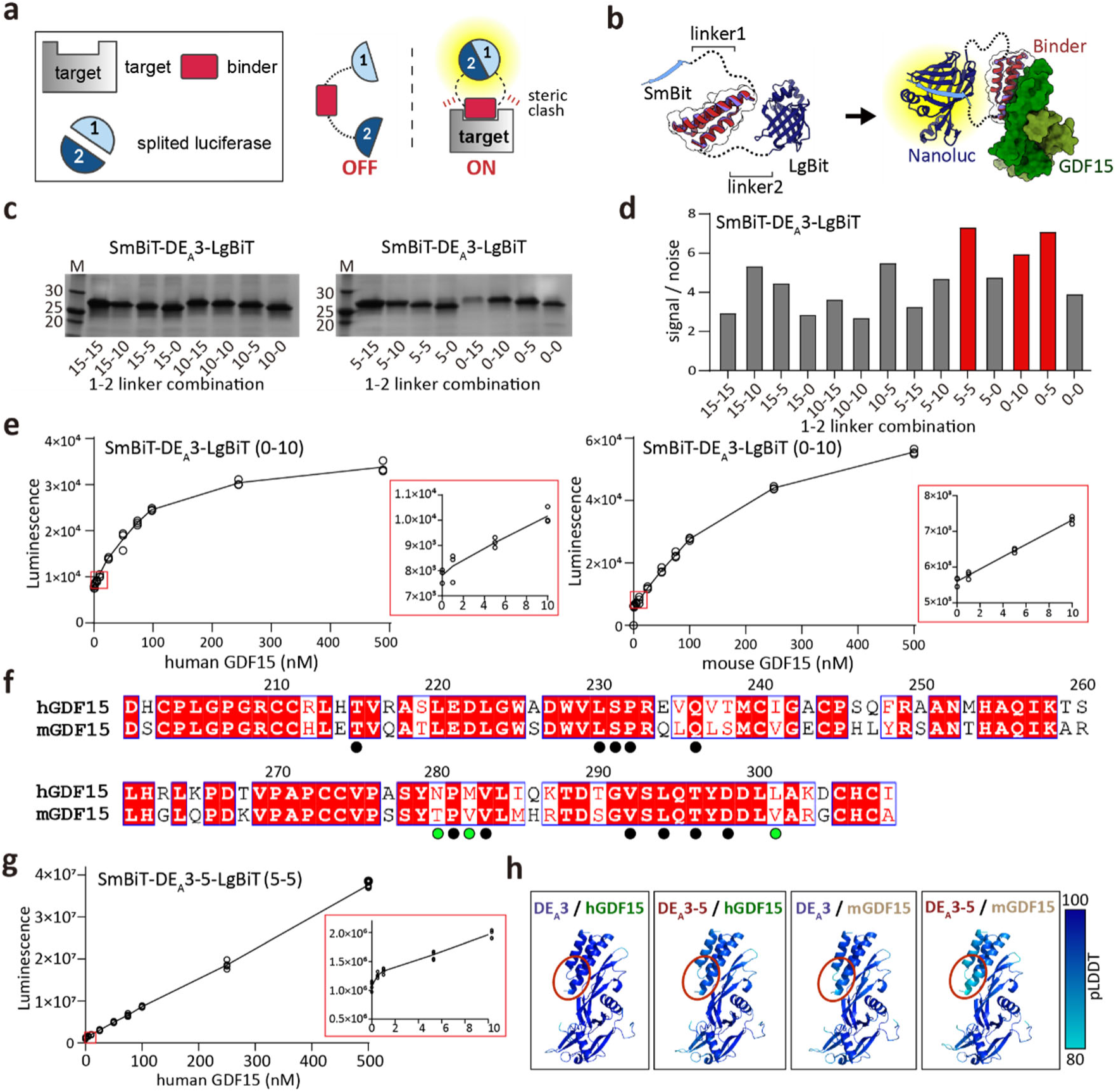
Development of a binding-activated tandem split-enzyme (BAT) biosensor for GDF15 detection. **a, b.** Schematic illustration (a) and structural model (b) of the BAT biosensor (SmBiT-GDF15 binder-LgBiT) for GDF15 detection in the “OFF” and “ON” states. The BAT biosensor consists of a designed GDF15 binder (red) flanked by SmBiT at the N-terminus (cyan, 1, SmBiT) and LgBiT at the C-terminus (blue, 2, LgBiT). In the absence of GDF15, two split luciferase fragments remain apart (“OFF” state with only background activity). Upon GDF15 binding, steric constraints bring two split luciferase fragments into proximity, enabling fragment complementation and restoring NanoLuc activity (“ON” state). The lengths of linker1 and linker2 (b, left) are key determinants for background signals in the absence of GDF15. **c.** SDS-PAGE analysis of SmBiT-DE_A_3-LgBiT with various linker combinations after *E.coli* expression and affinity purification. **d.** Screening of linker combinations for SmBiT-DE_A_3-LgBiT using a luminescence assay. Signal-to-noise ratios (luminescence intensity of each construct divided by that of the control without GDF15) are shown, with optimal linker combinations highlighted in red. **e, g.** Luminescent signals of SmBiT-DE_A_3-LgBiT with 0-10 linkers (e) and of SmBiT-DE_A_3-5-LgBiT with 5-5 linkers (g). Luminescence (arbitrary units, AU) is plotted against various concentrations of human or mouse GDF15 (n = 3). The linear detection range is indicated with a red box (0–10 nM). **f.** Sequence alignment of human and mouse GDF15. Conserved residues at site A are marked with black circles; species-specific substitutions at the interface are highlighted with green circles. **h.** AF3-predicted structures of DE_A_3 or DE_A_3-5 bound to human or mouse GDF15. Per-residue pLDDT values are color-coded according to the scale bar.

We evaluated BAT sensors constructed from four top-performing binders (SG_A_2, SG_A_2-4, DE_A_3, and DE_A_3-5), which are derived from distinct design strategies (Figs. 2a, 3a, and 4d). Because signal output depends on the spatial orientation of luciferase fragments, we systematically optimized linker lengths between the binder and each luciferase fragment (SmBiT-linker1-GDF15 Binder-linker2-LgBiT-His_6_). Four linker lengths (0, 5, 10, and 15 residues) were tested for both linker1 and linker2, generating 16 unique configurations per binder.

BAT sensors based on SG_A_2 and SG_A_2-4 exhibited modest signal-to-noise ratios (<4) across all linker combinations, likely due to the parallel orientation of their N- and C-termini, which restricts the steric displacement required for efficient luciferase complementation (Fig. S5). The most effective SmBiT-linker1-SG_A_2-linker2-LgBiT construct (10 a.a for linker 1 and 10 a.a for linker 2, 10-10 linker) could detect human GDF15 within the 10–100 nM concentration range, but not mouse GDF15 (Fig. S5d and S5e). Conversely, the SmBiT-DE_A_3-LgBiT biosensor, characterized by oppositely oriented N- and C-termini, yielded markedly higher signal-to-noise ratios. Among the tested constructs, those with 0–10, 0–5, and 5–5 linker lengths exhibited the best performance (Fig. 5c-d). During the expression and purification of SmBiT-DE_A_3-LgBiT, the construct with a 10–0 linker proved difficult to express, and the 0–15 linker construct tended to aggregate; consequently, both were excluded from subsequent testing. When assessed across a concentration range of 0–100 nM of human GDF15, the 0–10 construct demonstrated the most linear dose-response curve and demonstrated consistent reproducibility, thereby selecting for subsequent analysis (Fig. 5e, left). Similar results were observed for the 0–5 and 5–5 linker combinations against human GDF15 (Fig. S5i and S5j).

Sequence alignment of human and mouse GDF15 revealed 72% identity, with key hydrophobic residues at site A highly conserved (Fig. 5f). Consistent with this, the SmBiT-DE_A_3-LgBiT biosensor (0–10 linker) demonstrated a robust, concentration-dependent luminescence for both human and mouse GDF15, showing near-linear detection across 1–100 nM and maintaining performance in serum supplemented with recombinant GDF15 (Fig. 5e, right). These results underscore its potential for cross-species and clinically relevant detection.

Although structural predictions indicated that DE_A_3-5 possesses a scaffold highly similar to that of DE_A_3, the 0–10 linker construct for DE_A_3-5 could not be successfully expressed. Consequently, the biosensor incorporating the 5–5 linker configuration, which displayed superior detection performance among the alternative constructs, was utilized in subsequent experiments. Notably, the SmBiT-DE_A_3-5-LgBiT biosensor (5-5 linker) detected only human GDF15 with sub-nanomolar sensitivity, exhibiting a linear detection range of 0.5-500 nM (Fig. 5g). However, it failed to sensitively detect mouse GDF15 (Fig. S5k). This limitation likely arises from subtle sequence differences at the interaction interface (N280, M282, L301 in human GDF15 versus T280, V282, V301 in mouse GDF15) (Fig. 5f), which may alter the positioning of the DE_A_3-5 N-terminus. AF2 predictions of the DE_A_3-5/mouse GDF15 complex further support this interpretation, showing markedly reduced pLDDT values at the N-terminal interface compared to other complexes (DE_A_3/human GDF15, DE_A_3/mouse GDF15, and DE_A_3-5/human GDF15) (Fig. 5h, red circle), indicating increased flexibility that likely impaired luciferase complementation.

Collectively, our results demonstrate that BAT biosensors incorporating GDF15 binders provide a robust and versatile platform for rapid, cross-species detection of GDF15 with high sensitivity, underscoring their potential for clinical diagnostic applications.

### 2.6. Therapeutic potential of an Fc-fusion GDF15 binder

To assess the therapeutic potential of the highest-affinity binder, we generated an Fc-fusion GDF15 binder to inhibit GDF15/GFRAL/RET signaling axis. As an initial approach, a bispecific Fc-fusion construct was attempted by combining two designed binders targeting distinct sites on the GDF15 dimer (SG_A_2-4 for site A and SSG_B_2 for site B). However, this bispecific construct exhibited poor expression yields, precluding its further development (Fig. S6). Consequently, a homodimeric design was employed, wherein the high-affinity site A binder SG_A_2-4 was fused to both arms of a human IgG Fc domain (Fig. 6a). The resulting SG_A_2-4-Fc protein was robustly expressed in a mammalian expression system and purified to high homogeneity and purity (Fig. 6b).

**Figure 6.**
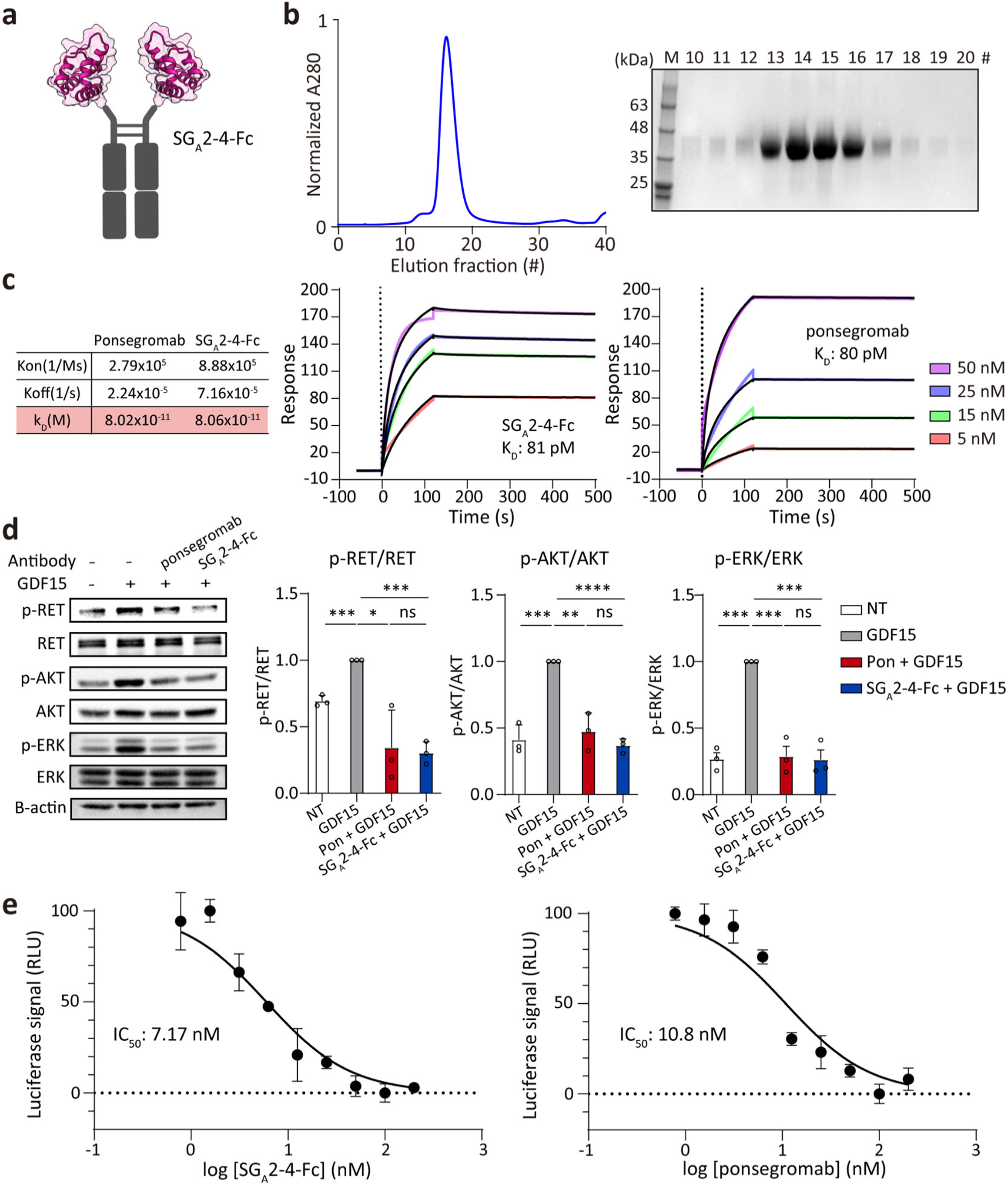
Development of GDF15 decoy receptors using GDF15 site A binder. a. Schematic diagram of Fc-fused SG_A_2-4 binder (SG_A_2-4-Fc). b. SEC profile of SG_A_2-4-Fc on a Superdex® 200 Increase 10/300 GL column (left) and SDS-PAGE analysis of elution fractions (right). c. SPR analysis of SG_A_2-4-Fc and ponsegromab binding to immobilized GDF15. Sensorgrams are shown for analytes ranging from 5 to 50 nM (SG_A_2-4-Fc or ponsegromab). d. Inhibition of GDF15-induced RET, AKT, and ERK phosphorylation in HEK293T cells stably expressing GFRAL and RET. Cells were co-treated with GDF15 (10 nM, 246 ng/ml) and either SG_A_2-4-Fc or ponsegromab (100 nM; 7.8 μg/ml for SG_A_2-4-Fc or 14.6 μg/ml for ponsegromab) for 30 mins. Phosphorylation was quantified relative to total protein (RET, AKT, and ERK each), normalized to the GDF15-only condition (n = 3). Statistical significance was determined using unpaired t-test (****P < 0.0001; ***P < 0.001; **P < 0.01; *P < 0.05; ns, not significant). e. Dose-dependent inhibition of GDF15-induced SRE-luciferase activity by SG_A_2-4-Fc or ponsegromab in HEK293T cells co-expressing GFRAL, RET, and an SRE-Luc2 reporter. Data were normalized to GDF15-induced luminescence (100%), and IC_50_ values were determined by non-linear regression fitting in GraphPad Prism.

Biophysical characterization revealed that Fc-mediated dimerization of SG_A_2-4 conferred a marked avidity gain, enhancing the apparent binding affinity from 330 pM for the monomeric binder to 81 pM for the Fc-fusion. Notably, this affinity is indistinguishable from that of ponsegromab (K_D_=80 pM), a clinical-stage anti-GDF15 antibody currently in Phase II trials for treating cancer cachexia^20^ (Fig. 6c). We next tested the functional activity of SG_A_2-4-Fc in blocking GDF15 signaling. In HEK293 cells stably expressing human GFRAL and RET, GDF15 stimulation induced phosphorylation of RET, AKT, and ERK. However, Co-treatment with either SG_A_2-4-Fc or ponsegromab (100 nM) significantly suppressed phosphorylation of all three proteins (Fig. 6d). In parallel SRE-luciferase reporter assays, SG_A_2-4-Fc inhibited GDF15-induced GFRAL/RET signaling in a dose-dependent manner (IC₅₀=7.2 nM), demonstrating a potency comparable to ponsegromab (IC₅₀=10.8 nM) under identical experimental conditions. (Fig. 6e).

Collectively, these results demonstrate that SG_A_2-4-Fc not only attains a binding affinity comparable to that of the clinical anti-GDF15 antibody ponsegromab but also effectively inhibits downstream GDF15 signaling. This highlights its potential as a promising alternative therapeutic approach, derived from a fully *de novo* designed protein scaffold, for the treatment of cancer cachexia.

## 3. Discussion

The development of protein binders exhibiting high affinity and specificity has long been a central challenge, with extensive implications for both diagnostic and therapeutic applications. Recent advancements in artificial intelligence—such as AlphaFold for structural prediction, RFdiffusion for backbone generation, and ProteinMPNN for sequence design—are now enabling *de novo* protein design with customized folds and functions^33,34^. This progression signifies a paradigm shift beyond traditional protein engineering approaches^24,26,35–38^. Nonetheless, significant limitations persist in *de novo* binder design, especially for targets characterized by flat or polar surfaces, where shape complementarity is constrained, success rates remain modest, and extensive experimental screening is required^39,40^. Here, we addressed these challenges through natural scaffold–based design informed by evolutionary complexes, and benchmarked this strategy against diffusion-based strategies, ultimately generating high-affinity binders against GDF15, a key driver of cancer cachexia. These binders enabled the development of both a highly sensitive GDF15 biosensor and a potent Fc-fused decoy receptor, underscoring their potential to address the unmet clinical need in cancer cachexia.

First, we systematically evaluated three complementary strategies for generating initial scaffolds to design GDF15 binders: (1) scaffold grafting (SG), (2) diffusion-based *de novo* design, and (3) scaffold-search and grafting (SSG). These were applied to distinct GDF15 epitopes, either the convex finger-loop surface (site A) or the opposing shallow concave surface (site B). The SG strategy transplanted a helix– loop–helix motif from the native GFRAL D2 domain, which naturally engages GDF15 site A. This pre-organized geometry ensured tight shape complementarity^38,41–43^, the simple sequence optimization with ProteinMPNN resulted in stable binders that maintained the triangular scaffold architecture and improved affinity from approximately 90 nM in the parental segment to single-digit nanomolar levels. Of note, nearly 30% of variants satisfied *in silico* filtering metrics at the initial sequence-design step (28 out of 100 designs, pLDDT > 85, pAE_interaction < 10, ΔΔG < –30). More significantly, a single iteration of subsequent scaffold-guided partial diffusion and sequence redesign proved sufficient to enhance binding affinity by an order of magnitude, resulting in sub-nanomolar affinities and approximately a 300-fold increase in affinity relative to the native GFRAL D2 scaffold (GFRAL D2 K_D_ = 90 nM; SG_A_2 K_D_ = 5 nM; SG_A_2-4 K_D_ = 300 pM). While this efficiency highlights the robustness of grafting pre-formed interfaces, all SG-derived binders maintain parallel N- and C-terminal orientations analogous to GFRAL D2, thereby limiting their applicability in sensitive BAT-based biosensor platforms.

In contrast, the diffusion-based *de novo* approach explored entirely novel topologies unconstrained by natural templates. Although the resulting designs, guided only by hotspot constraints and amino acid length (exposed hydrophobic residues on GDF15 site A, 50–90 residues), mostly adopted three-helix bundles, these differed markedly from SG-derived scaffolds. Among the *in silico* filtered candidates (10.4%, 539 out of 5,184 designs, pLDDT > 85, pAE_interaction < 10, ΔΔG < –30), we experimentally validated the top five, identifying soluble binders with low nanomolar affinity in the initial design round. These findings underscore the efficacy of generative *de novo* design via RFdiffusion. However, further refinement through partial diffusion did not significantly enhance the affinity of the parental binder DE_A_3 (DE_A_3 K_D_ = 7.5 nM and DE_A_3-5 K_D_ = 6.9 nM), and this strategy also failed to generate high-affinity binders for the polar-rich GDF15 site B. The development of entirely novel backbones significantly broadens the search space, thereby facilitating the identification of new topologies for complex epitopes. However, this approach is computationally demanding for scaffold generation, sequence design, large-scale structural prediction, and *in silico* filtering.

The scaffold-search and grafting (SSG) strategy proved to be essential for the binder design of GDF15 site B, a notably challenging epitope due to its shallow topology and sparse hydrophobic patches. While scaffold-guided and *de novo* strategies were unsuccessful in producing reliable binders at this site, SSG addressed these limitations by mining evolutionary structural analogs from the PDB. A DALI search identified Repulsive Guidance Molecule A (RGMA), which naturally binds BMP2—a TGF-β family cytokine structurally homologous to GDF15—as a suitable initial scaffold. By structurally aligning GDF15 in place of BMP2 within the RGMA complex, followed by partial diffusion and sequence design, we generated binder candidates that retained RGMA’s stable α-helical bundle while successfully adapting to the GDF15 surface. Although only 0.3% of the sequence-designs passed *in silico* filtering (10 out of 3,000 designs; pLDDT > 85, pAE_interaction < 10, ΔΔG < –30), four of the five designs folded stably in E. coli, with one exhibiting measurable binding (SSG_B_2 K_D_ = 121 nM). These findings underscore the significance of repurposing natural complexes as structural blueprints to tackle design challenges at otherwise intractable epitopes. Furthermore, these insights are likely applicable across the TGF-β superfamily, which shares a conserved cystine-knot growth factor (CKGF) domain topology characterized by modular finger loops that facilitate receptor binding interactions^44^. Our methodology, notably SSG, provides a framework for targeting comparably challenging interfaces within other members of the TGF-β superfamily, such as TGF-β1, Activin A, and BMP9, thereby expanding opportunities for the development of both therapeutic antagonists and diagnostic biosensors. Taken together, these three strategies—SG for rapid optimization, *de novo* diffusion for topological exploration, and SSG for challenging surfaces—constitute a robust and versatile pipeline for designing high-affinity binders.

An immediate and compelling application of our *de novo* GDF15 binders lies in the field of diagnostics, particularly where circulating GDF15 serves as a crucial pathophysiological marker. Elevated GDF15 levels are associated with cancer cachexia, chemotherapy-induced nausea, and pregnancy-related hyperemesis gravidarum^37,38,39^, among other conditions in which early detection could inform timely intervention before clinical decline. While FDA-approved immunoassays, such as Roche’s Elecsys® offer precise quantification, they require centralized laboratory infrastructure and multi-step procedures, thereby restricting their applicability in point-of-care or low-resource environments. To address this limitation, we developed a binding-activated tandem split-enzyme (BAT) biosensor^23^ by fusing our high-affinity GDF15 binder to complementary fragments of NanoLuc luciferase. Upon GDF15 engagement, the sensor reconstitutes luciferase activity, emitting a luminescent signal in a single-step, wash-free assay. Through systematic optimization of linker lengths and domain geometries, the sensor achieved robust detection across the clinically relevant range (sub-nanomolar to 100 nM). Notably, it functions reliably in both buffer and serum, exhibiting cross-species reactivity that facilitates its application in both preclinical models and human samples. Furthermore, as demonstrated by Ravalin et al., lateral flow assay (LFA) adaptation of similar luciferase-based systems may pave the way for low-cost, user-friendly diagnostic formats^32^. As the repertoire of AI-designed binders expands, we envision a rapid development of analogous biosensors for a broad spectrum of disease markers, metabolic signals, and infectious pathogens. However, current BAT biosensors still require extensive empirical linker optimization. Moving forward, predictive and structure-informed frameworks for the optimization of BAT’s linker will be essential for generalizing and expediting the rational design of robust, modular biosensors based on the binder-fused BAT platform.

In parallel with diagnostic developments, our high-affinity GDF15 binders were engineered into Fc-fusion proteins to serve as therapeutic decoy receptors. This format dimerizes via the Fc domain, thereby enhancing apparent affinity from 300 pM (SG_A_2-4) to 81 pM (SG_A_2-4-Fc) through avidity. The Fc-fused GDF15 binder exhibited neutralization potency comparable to Pfizer’s anti-GDF15 monoclonal antibody, ponsegromab, in cell-based assays, despite being developed entirely *in silico* without animal immunization or phage library screening. While we have not yet evaluated pharmacokinetics, the Fc-fused binder is expected to prolong serum half-life through neonatal Fc receptor recycling, as evidenced by other Fc-based therapeutics^45–47^. The modularity of *de novo* binders can also be leveraged for the rapid generation of bispecific or multifunctional agents. For instance, dual-target Fc constructs could be engineered to concurrently inhibit GDF15 and other disease-associated factors, thereby enabling tailored interventions for multifactorial diseases. Unlike traditional antibodies, such constructs can be rationally designed and optimized through iterative *in silico* workflows. Although immunogenicity remains a pertinent consideration for non-natural proteins, prior studies suggest that hyperstable mini-proteins may exhibit low immunogenicity due to their resistance to proteolytic degradation and antigen presentation^48–50^. Furthermore, potential T-cell epitopes can be computationally predicted and removed without compromising structure or function. Altogether, these results highlight the potential of computationally designed binders as a flexible and rapidly deployable platform for therapeutic development^51,52^.

In summary, our complementary binder design pipeline, encompassing scaffold grafting, *de novo* backbone generation, and scaffold-search & grafting, delineates how structure-based intuition and AI-driven design can be strategically integrated. Each approach offers distinct strengths, and their synergy enables the generation of potent and versatile binders targeting structurally challenging epitopes. As generative AI models and predictive technologies continue to advance, such frameworks for binder design will accelerate the development of next-generation diagnostic and therapeutic proteins.

## 4. Experimental Section/Methods

### Three complementary scaffold design strategies for GDF15 binder

To generate antagonistic binders targeting site A of GDF15, we first employed a scaffold grafting (SG) strategy with the D2 domain of GFRAL as the structural template (PDB: 6Q2J)^23^. The initial scaffold was created by extracting the triangle-shaped helix–loop–helix motif that directly engages GDF15, and 100 sequences for a scaffold were generated using the ProteinMPNN-FastRelax protocol described in Bennett et al without residue restrictions. This protocol begins with a round of ProteinMPNN, followed by iterative cycles between FastRelax and ProteinMPNN to improve the sequence-structure agreement. Sequence sampling was conducted with a temperature parameter (T = 0.0001) to ensure sequence diversity. Complex structures of the designed sequences with GDF15 were predicted using AlphaFold3^26^, and the designs were ranked and filtered based on ΔΔG (ΔΔG ≤ –30 kcal/mol), predicted aligned error (pAE_interaction <10), and predicted Local Distance Difference Test (pLDDT > 85, per-residue confidence scores). The two designs with the best scores were selected, considering Rosetta scores after FastRelax for experimental validation^53^.

For diffusion-based *de novo* design, we used RFdiffusion (T=50) to generate each backbone without relying on pre-existing scaffolds. Backbone lengths were constrained to 50–90 residues, and hydrophobic residues (L230, V283, I285, V292, and L294 for site A; M253, W285, W288, and Y297 for site B) were specified as hotspot constraints to guide interface formation with GDF15. Each backbone was subsequently sequence-designed as described above. Designs were evaluated using AlphaFold2 (AF2) using an initial guess and target templating, and candidates with pLDDT values > 85, pAE_interaction < 10, ΔΔG ≤ –30 kcal/mol, and a favorable interface geometry were selected for experimental characterization.

To design binders against the structurally distinct site B of GDF15, we implemented a scaffold-search and grafting (SSG) strategy. The Dali server^54^ was used to query the GDF15 dimer against the Protein Data Bank to identify structurally similar proteins. Apo-form structures resembling the GDF15 dimer alone were excluded, and only PDB entries in which GDF15-like folds were engaged in protein– protein interactions were retained. For each, the GDF15 dimer was structurally superimposed onto GDF15-like domains, which were then replaced while preserving the original interacting partner. Partial diffusion was subsequently performed to generate interface-compatible backbones with the GDF15 dimer. Among these, the scaffold that preserved its fold without collapsing after partial diffusion was chosen as the optimal SSG template. From this analysis, we identified a scaffold derived from the Repulsive Guidance Molecule A (RGMA) (PDB codes: 4UHY), which naturally interacts with BMP2, cytokines structurally homologous to GDF15.

To optimize and diversify experimentally validated binders (SG_A_2 and DE_A_3) and the RGMA-derived scaffold, we applied RFdiffusion in a partial diffusion mode, which allows the input structures to be noised only up to a user-specified timestep rather than undergoing full noising^39,55^. Input scaffolds were superimposed with the GDF15 dimer at the target site (site A or site B). Hydrophobic residues at site A (L230, V283, I285, V292, and L294) or site B (M253, W285, W288, and Y297) were designated as hotspot constraints (T=0∼25). The resulting backbones were sequence-designed using the ProteinMPNN-FastRelax protocol (10 sequences per backbone) and evaluated with AF2 initial guess and template-based modeling.

Final candidates were selected based on pLDDT > 85, pAE_interaction < 10, ΔΔG ≤ –30 kcal/mol, and a favorable interface geometry for experimental validation.

### Cell lines and cell culture

Expi293F cells (#A14527, Thermo Fisher Scientific) were maintained in Expi293 Expression Medium (#A1435102, Thermo Fisher Scientific) at 37°C in a humidified incubator with 8% CO₂ under constant shaking. The HEK293 reporter cell line, which stably expresses human GFRAL, human RET, and an SRE-Luc2 luciferase reporter gene (293T-SRE384-Luc2-RET-GFRAL; #KC-2134, Kyinno Biotechnology) was cultured in Dulbecco’s Modified Eagle Medium (DMEM) supplemented with 10% fetal bovine serum (FBS; #12483020, Gibco), 1 µg/ml puromycin (#P8833, Sigma-Aldrich), and 150 µg/ml hygromycin B (#10687010, Gibco).

### Construct, expression, and purification of recombinant proteins

All cloning was performed using the EZ-Fusion™ HT Cloning Kit (#EZ015TM, Enzynomics). The gene encoding mature human GDF15 (residues 198–308; GenBank accession no. 9518) was inserted into the *NotI* and *XhoI* sites of a modified pcDNA3.4 vector (#A14697, Invitrogen) containing an N-terminal Twin-Strep tag followed by a thrombin cleavage sequence. For expression of the mature GDF15 dimer, an N-glyco-engineered variant (R198N and G200T) with improved expression yield and preserved biological activity was used^56^. The gene encoding hGFRAL D2 (residues 115–217) was subcloned into a modified pcDNA3.1 mammalian expression vector (#V79020, Invitrogen) containing a thrombin protease cleavage site and C-terminal Fc tag. Genes encoding the designed GDF15 binders (SG_A_1∼SG_A_2, SG_A_2-1∼SG_A_2-5, DE_A_1∼DE_A_5, DE_A_3-1∼DE_A_3-7, and SSG_B_1∼SSG_B_5) were synthesized by Twist Bioscience and cloned into a pPROEX-HTa *E. coli* expression vector containing an N-terminal 6×His tag and a TEV protease cleavage site.

For SmBiT-Binder-LgBiT-His_6_ constructs, the SmBit (residues 1∼11)–linker1–binder–linker2–LgBit (residues 12∼171), synthesized by Twist Bioscience, was cloned into the pPROEX-HTa vector using EcoRI and XhoI restriction sites. Linker1 and linker2 were 0, 5, 10, or 15 amino acids in length, generating 16 combinations per binder. For SG_A_2-4-Fc, the gene encoding SG_A_2-4 was subcloned into a modified pcDNA3.1 mammalian expression vector (#V79020, Invitrogen) containing a C-terminal Fc tag, thrombin protease cleavage site, and 6x His tag.

For *E. coli* expression, recombinant plasmids encoding GDF15 binder–His₆, or SmBiT-Binder-LgBiT-His_6_ were transformed into *E. coli* BL21 (DE3) and grown in LB medium containing 100 μg/ml ampicillin at 37°C. At an OD₆₀₀ ≈ 1.0, protein expression was induced with 0.5 mM IPTG, followed by incubation at 30°C for 6 h. Cells were harvested by centrifugation and resuspended in a lysis buffer (20 mM Tris–HCl, pH 8.0, 150 mM NaCl). Lysis was performed by sonication on ice, and cell debris was removed by centrifugation at 15,000 × g for 30 min at 4°C. The supernatant was loaded onto Ni–NTA agarose resin (#30210, Qiagen) pre-equilibrated with a lysis buffer. Bound proteins were then eluted with a lysis buffer supplemented with 250 mM imidazole. Eluted proteins were further purified by size-exclusion chromatography (SEC) on a Superdex® 200 Increase 10/300 GL column (#GE28-9909-44, Cytiva) equilibrated with a lysis buffer. Fractions corresponding to the major peaks were pooled, concentrated to 3 mg/ml, aliquoted, and stored at −80°C.

For mammalian expression, plasmids encoding the Twin-Strep-GDF15 dimer, SG_A_2-4–Fc-6xHis, or hGFRAL D2-Fc (1 μg) were transfected into Expi293F cells (#A14527, Thermo Fisher Scientific; 2 × 10⁶ cells/ml) using the ExpiFectamine Transfection Kit (#A14524, Thermo Fisher Scientific). Cells were cultured at 37 °C under 8% CO₂, shaking at 125 rpm for 5 days. Expression enhancers were added 18–22 h post-transfection. Supernatants containing recombinant GDF15 dimer were clarified by centrifugation and loaded onto Strep-Tactin XT Sepharose resin (#2-1201-025, IBA Life Sciences). Bound proteins were washed with four column volumes of wash buffer (#2-1002-001, IBA Life Sciences; 20 mM Tris–HCl, pH 8.0, 150 mM NaCl, 1 mM EDTA) and eluted with a buffer containing 5 mM desthiobiotin. Eluted GDF15 was concentrated to 5 mg/ml using Amicon Ultra centrifugal filters (#UFC8030, Millipore) and further purified by SEC on a Superdex® 200 Increase 10/300 GL column (#GE28-9909-44, Cytiva) equilibrated in 20 mM Tris–HCl, pH 8.0, and 150 mM NaCl buffer. For SG_A_2-4-Fc-6xHis purification, the supernatants were applied to Ni-NTA (# 30230, Qiagen) and washed with 10 column volumes (CV) of wash buffer (20 mM Tris-HCl, pH 8.0, 200 mM NaCl, 20 mM imidazole). For purification of hGFRAL D2, the supernatants containing hGFRAL D2-Fc were loaded onto Protein A resin (#1010200, Amicogen) and washed with 10 column volumes (CV) of PBS. In both cases, bound proteins were digested on-resin overnight at 4°C with thrombin (enzyme-to-substrate ratio 1:100) in PBS. The cleaved proteins (SG_A_2-4-Fc or hGFRAL D2) were further purified by SEC on a Superdex 200 Increase 10/300 GL column (Cytiva) equilibrated with PBS.

### Yeast surface display

Genes encoding the top 100 DE_A_3-guided partial diffusion binder candidates were synthesized (Twist Bioscience) and cloned into the pETCON vector for yeast surface display. The plasmids were transformed into *Saccharomyces cerevisiae* strain EBY100 using the lithium acetate/PEG method (#T2001, Zymo Research). Transformants were selected on SDCAA agar plates at 30°C for 48 h, then transferred into 3 mL SDCAA medium (#2S0540, Teknova) and cultured at 30°C for 48 h until reaching OD₆₀₀ ≈ 1.0. For the induction of designed binder candidates on the yeast surface, cells were centrifuged (2,000 g, 10 min, 4°C) and resuspended in SGCAA medium (#2S0542, Teknova) at a concentration of 1.4 × 10⁷ cells/mL, followed by incubation at 30°C for 36 h. Induced cells were centrifuged (2,000 g, 5 min), washed once with ice-cold PBSA (PBS with 1% w/v BSA), and resuspended in PBSA for labeling.

To assess binder surface display, 100 μL of cells were incubated with anti-Myc-PE-Cy7 antibody (1:50 dilution, 3 μg/mL; #36271, Cell Signaling Technology) for 30 min at room temperature, then washed twice with PBSA to remove excess unbound antibody. For GDF15 binding assessment, cells were subsequently incubated with twin-strep–tagged GDF15 (1 μM) and strep-tactin-PE (1:1,000 dilution, 75 μg/mL) in PBSA for 30 min at room temperature, followed by three times washing with ice-cold PBS.

Fluorescence-activated cell sorting (FACS) was performed using a CytoFLEX SRT cell sorter (#B75489, Beckman Coulter) with gating optimized for double-positive selection (Myc with PE-Cy7 and strep-tag with PE). Data was analyzed using CytExpert SRT software (Beckman Coulter). Approximately 3,000 double-positive cells were sorted into 5 mL SDCAA recovery medium and cultured at 30°C for 36 h.

Cells were then plated on SDCAA agar and grown at 30°C for 48 h. Forty individual colonies were picked, plasmids extracted (Zymoprep Yeast Plasmid Miniprep II, #D2004, Zymo Research), and each clone sequenced by Sanger sequencing, identifying seven unique binder candidates.

### Binding kinetics between GDF15 binders and recombinant GDF15 dimer

Binding affinities of GDF15 binders, binder–Fc, and ponsegromab against recombinant GDF15 dimers were measured by surface plasmon resonance (SPR) using a Biacore T200 system equipped with CM5 Series S sensor chips (#BR100399, GE Healthcare). Recombinant GDF15 dimers were immobilized on CM5 chips (#BR100530, Cytiva) by amine coupling in 10 mM sodium acetate, pH 5.5.HBS-EP+ buffer (0.01 M HEPES, pH 7.4, 0.15 M NaCl, 3 mM EDTA, 0.05% v/v surfactant P20; #BR100669, Cytiva) was used as both running and sample buffer.

Analytes (GDF15 binders, SG_A_1, SG_A_2, SG_A_2-1, SG_A_2-3, SG_A_2-4, SG_A_2-5, DE_A_3, DE_A_4, DE_A_5, DE_A_3-1, DE_A_3-2, DE_A_3-5, SSG_B_1, SSG_B_2, SSG_B_3, and SSG_B_5; SG_A_2-4-Fc or Ponsegromab) were diluted in HBS-EP+ buffer and injected over the GDF15-immobilized sensor chips at 0-100 nM (5, 15, 25, 50, and 100 nM) for 300 s at a flow rate of 30 μl/min, followed by dissociation for 300 s in the running buffer. The chip surfaces were regenerated after each cycle with 10 mM glycine (pH 2.0). Association (kon, M⁻¹s⁻¹) and dissociation (koff, s⁻¹) rate constants were determined over 300-second intervals, and the equilibrium dissociation constant (K_D_, M) was calculated as koff/kon. Kinetic parameters were obtained using Biacore Insight Evaluation Software: a two-state reaction model was applied to the minibinders, and a 1:1 binding model was used for SG_A_2-4-Fc and Ponsegromab.

### Luminescence-based BAT biosensor assay for GDF15 quantification

Recombinant human or mouse GDF15 (#9279-GD or #8944-GD, R&D Systems) was prepared at a stock concentration of 250 μg/mL and subsequently diluted in pooled human serum (Sigma #H6914) or mouse serum (Abbkine #BMS0070), resulting in a final concentration of up to 5 μg/mL (≤500 nM). The dilution was performed to ensure that 25 μL of the sample mixture contained no less than 97.5% serum (either human or mouse). The serum was used without prior heat inactivation and maintained at physiological pH (∼7.4). Sensor proteins (SmBiT-GF15 Binder-LgBiT-His6; stock concentration of 2.5 mg/mL) were diluted 1000-fold with 1× PBS (phosphate-buffered saline, pH 7.4, 137 mM NaCl, 10 mM phosphate, 2.7 mM KCl). Subsequently, 25 μL of the diluted biosensor solution was added to each well.

Each assay was conducted in a white, opaque 96-well plate (#3917, Corning), using a total reaction volume of 100 μL per well. The assay mixture consisted of 25 μL of serum-diluted GDF15 sample, 25 μL of PBS-diluted sensor protein, and 50 μL of Nano-Glo luciferase assay reagent (#N1110, Promega). After mixing serum-diluted GDF15 sample with PBS-diluted sensor protein, the plates were incubated at 4°C for 5 minutes, then gently mixed. Subsequently, the Nano-Glo luciferase assay reagent was added, and the plates were incubated for an additional 5 minutes at room temperature. Luminescence was subsequently measured at 560 nm using a SpectraMax iD5 plate reader (Molecular Devices).

### Signaling inhibition assays for the GDF15/GFRAL/RET pathway

Signaling inhibition of the GDF15/GFRAL/RET pathway was assessed using 293T-SRE-Luc2-RET-GFRAL cells (#KC-2134, KYinno), which stably expresses full-length human GFRAL, full-length human RET, and a firefly luciferase reporter gene under the control of serum response element (SRE).

For luciferase reporter assays, 293T-SRE-Luc2-RET-GFRAL cells (2.5 × 10⁴ cells/well) were seeded in 96-well plates with 100 μl of complete medium (high-glucose DMEM supplemented with 10% FBS, 1 μg/ml puromycin, and 150 μg/ml hygromycin B) and cultured overnight at 37°C. The next day, recombinant human GDF15 (10 nM) was added, along with serial dilutions of SG_A_2-4-Fc or ponsegromab (0.39–200 nM), and the mixture was incubated for 8 h at 37°C. Cells were lysed with 100 μl of Glo-lysis buffer (#E2661, Promega) for 10 min, followed by the addition of Bright-Glo luciferase reagent (#E2620, Promega) in a white opaque 96-well plate (#3917, Corning). After incubation for 5 min, luminescence was measured at 560 nm using a SpectraMax iD5 plate reader (Molecular Devices).

For western blotting, 293T-SRE-Luc2-RET-GFRAL cells (4 × 10⁵ cells/mL) were seeded in 6-well plates in complete medium and grown at 37°C to ∼80% confluence. Cells were treated with recombinant human GDF15 (10 nM; #9279-GD, R&D Systems) in the presence or absence of SG_A_2-4-Fc or ponsegromab (100 nM) for 30 min. After washing with cold PBS, cells were lysed in RIPA buffer (#RC2002-050-00, Biosesang) supplemented with phosphatase inhibitor (#4906845001, Roche) and protease inhibitor cocktail (#11836170001, Roche) for 60 min at 4°C. Lysates (10 μg protein per lane) were separated by SDS–PAGE and transferred to nitrocellulose membranes (#HATF00010, Millipore). Membranes were blocked with 5% skim milk in TBST (TBS with 0.1% Tween-20) and incubated overnight at 4°C with primary antibodies: anti-ERK (1:1000, #9102, Cell Signaling Technology), anti-p-ERK (1:2000, #4370, CST), anti-AKT (1:1000, #9272, CST), anti-p-AKT (1:2000, #4060, CST), anti-RET (1:1000, #14556, CST), anti-p-RET (1:500, #ab51103, Abcam), and anti-β-actin (1:1000, #sc-47778, Santa Cruz Biotechnology). After washing with TBS, membranes were incubated with HRP-conjugated secondary antibodies [anti-rabbit (#7074S, Cell Signaling Technology) for anti-ERK, anti-p-ERK, anti-AKT, anti-p-AKT, anti-RET, and anti-p-RET; anti-mouse (#62-6520, Invitrogen) for anti-β-actin] for 1 h at room temperature. Signals were visualized using enhanced chemiluminescence (#2332632, ATTO) and imaged on an iBright FL1500 system (Invitrogen). Band intensities were quantified using ImageJ (v1.54) and analyzed in GraphPad Prism (v8.0.2).

### Sequence alignment

Amino acid sequence alignments were performed using Clustal Omega with default parameters and visualized with ESPript 3.0. Fully conserved residues were indicated with red boxes, while residues with > 70% similarity in physicochemical properties were highlighted with blue boxes. Predicted secondary structures were annotated above the alignment, with α-helix shown as coiled symbols.

### Statistical analysis

All quantitative data are presented as mean ± Standard Error of the Mean (SEM), unless otherwise indicated. The number of biological and technical replicates (n) is specified in the corresponding figure legend, with n denoting the number of biological replicates. For immunoblotting, data are reported as mean ± Standard Deviation (SD). Data quantification was performed using ImageJ, and statistical analyses were conducted with GraphPad Prism v10.3.1. Statistical significance was assessed using an unpaired t-test, with significance thresholds defined as follows: P* < 0.05, P** < 0.01, P*** < 0.001, and P**** < 0.0001; ns indicates not significant.

## Acknowledgements

We are grateful to the staff of the Research Solution Center at IBS. Computational work for this research was performed using the high-performance computing resources in the IBS Research Solution Center. This work was supported by the National Research Foundation of Korea (RS-2024-00397681 to H.M.K. and RS-2025-00523575 to J.A.), the InnoCORE program of the Ministry of Science and ICT (N10250153 to H.M.K.), and the KAIST Convergence Research Institute Operation Program. We appreciate the careful reading and critical comments provided by Prof. Minho Shong (Graduate School of Medical Science & Technology, KAIST, Korea).

## Contributions

J.A. and H.M.K. conceived the overall study, designed the proteins and experiments, and analyzed the data. J.A., R.C., D.L., and S.K. generated recombinant constructs and performed protein expression and purification. J.A. conducted surface plasmon resonance (SPR) measurements, luminescence-based sensor assays, and yeast surface display screening. J.A. and S.K. carried out the GFRAL/RET ERK luciferase reporter assays. J.A., R.C., S.K., and H.M.K. wrote the manuscript.

## Notes

### Competing Interest Statement

The authors have declared no competing interest.

