## Supplementary information for "*De novo* and scaffold-based design of GDF15 binders for cancer cachexia diagnostics and therapeutics"

**This PDF file includes :**

Supplementary Figures S1 to S6

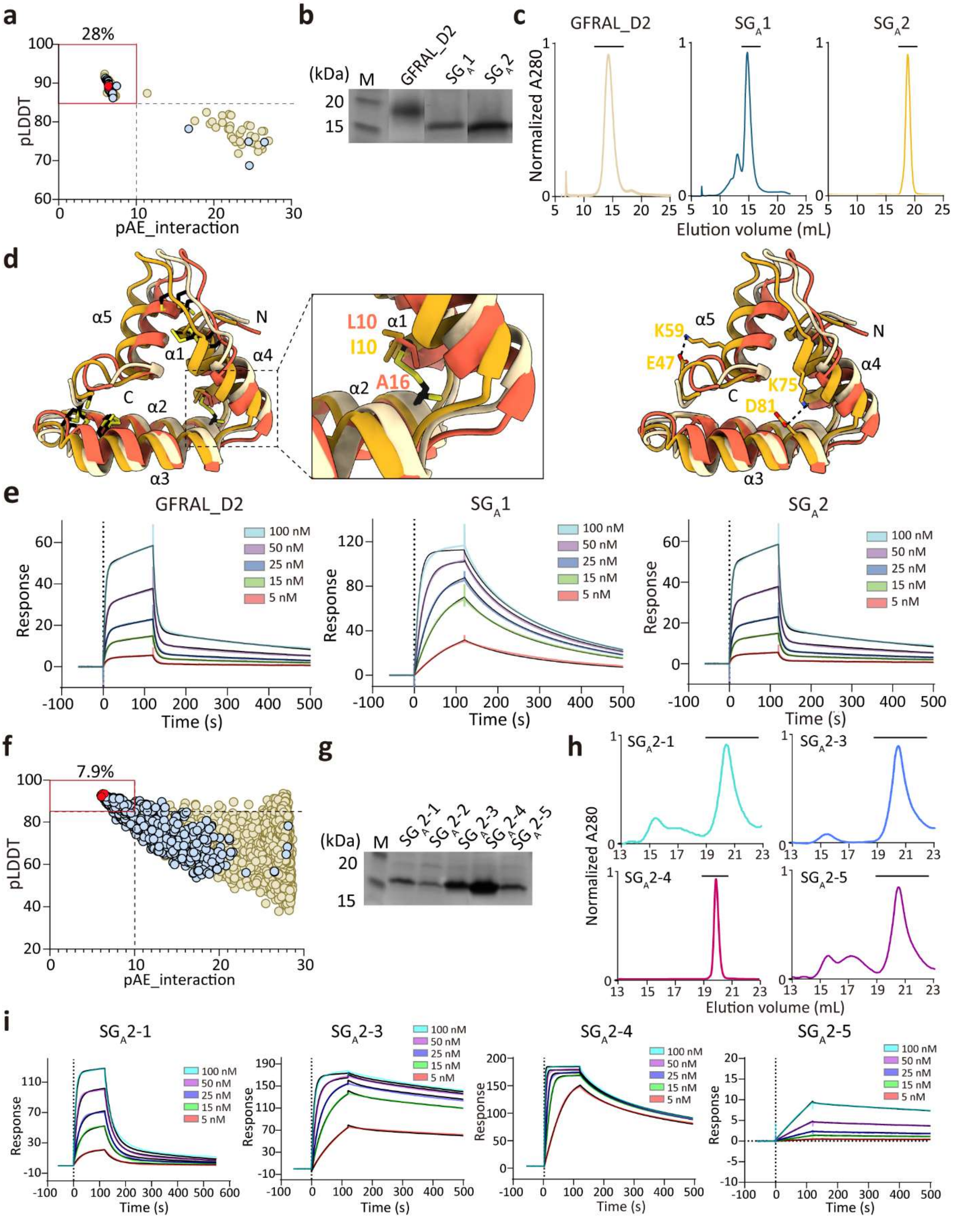


**Figure S1. Structural and biochemical characterization of scaffold-grafted GDF15 site A binders.**

**a, f.** Filtering metrics for initial GFRAL D2–based sequence designs **(a)** and scaffold-guided partial diffusion variants **(f)**. Candidates meeting the criteria of pLDDT > 85, pAE_interaction < 10, and ΔΔG < –30 are enclosed in red boxes. Overall, 28% of designs (28%, 28/100) from the first sequence design **(a)** and 7.9% from the partial diffusion designs (7.9%, 306/3850) **(f)** passed the filtering criteria. Data points with ΔΔG < –30 are shown in black; those with ΔΔG > –30 are shown in yellow. Among these, final candidates selected for experimental validation (protein expression, purification, and binding analysis) are indicated with red circles.

**b, g**. SDS-PAGE analysis of purified binders: **(b)** $SG_A1$ and $SG_A2$; **(g)** $SG_A2$-1 to $SG_A2$-5, after *E. coli* expression and affinity purification.

**c, h.** SEC profiles of designed binders: **(c)** GFRAL D2, $SG_A1$, and $SG_A2$; **(h)** $SG_A2$-1, $SG_A2$-3, $SG_A2$-4, and $SG_A2$-5.

**d.** Structural alignment of GFRAL D2 (beige), $SG_A1$ (orange), and $SG_A2$ (yellow), showing overall fold similarity with subtle structural differences. Residue substitutions at C10 and C16 (C10L/C16A in $SG_A1$ and C10I in $SG_A2$) disrupt the disulfide bond (left). In $SG_A2$, additional substitutions create new interactions between E47–K59 and K75–D81 that shift α3 and α5 (right).

**e, i.** SPR analysis of binding to recombinant GDF15 dimer: GFRAL D2, $SG_A1$, and $SG_A2$ **(e)**; $SG_A2$-1, $SG_A2$-3, $SG_A2$-4, and $SG_A2$-5 **(i)**. Sensorgrams are shown for analytes ranging from 5 to 100 nM (indicated binder).

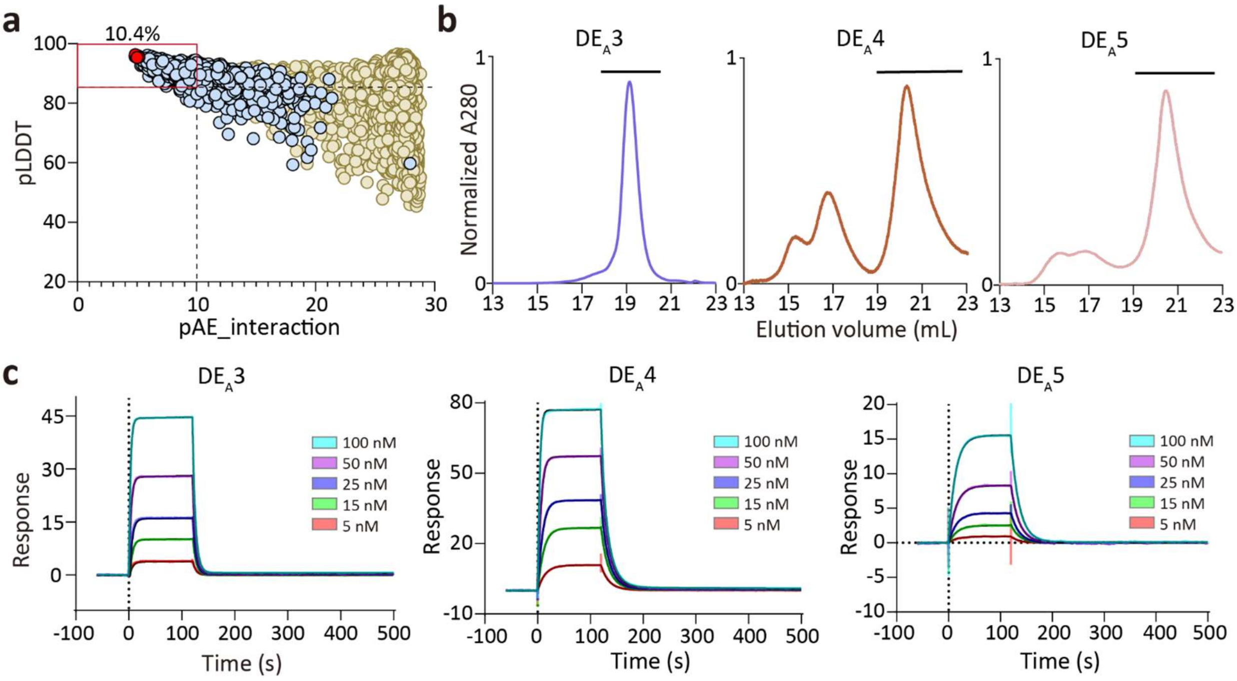


**Figure S2. Structural and biochemical characterization of *de novo* designed GDF15 site A binders.**

**a.** Filtering metrics for *de novo*–designed binders using RFdiffusion and ProteinMPNN. Candidates with pLDDT > 85, pAE_interaction < 10, and ΔΔG < –30 were selected (red box). Overall, 10.4% of the total designs (10.4%, 539/5,184) passed the filtering criteria. Data points with ΔΔG < –30 are shown in black; those with ΔΔG > –30 are shown in yellow. Among these, final candidates selected for experimental validation (protein expression, purification, and binding analysis) are indicated with red circles.

**b.** SEC profiles of *de novo* designed binders ($DE_A3$ to $DE_A5$).

**c.** SPR analysis of $DE_A3$, $DE_A4$, and $DE_A5$ binding to recombinant GDF15 dimer. Sensorgrams are shown for analytes ranging from 5 to 100 nM (indicated binder).

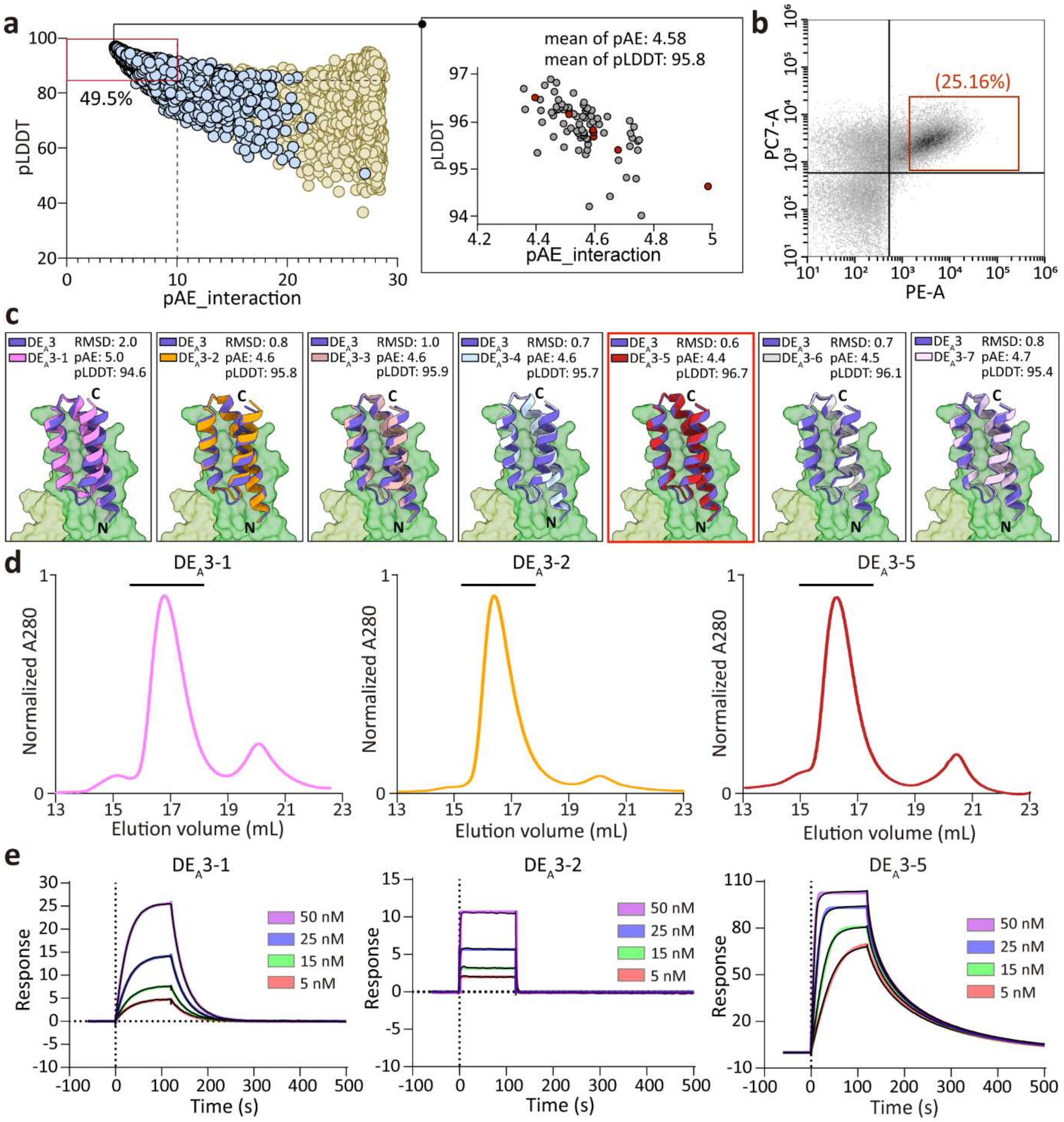


**Figure S3. Structural and biochemical characterization of DE$_A$3-derived GDF15 site A binders.**

**a.** Filtering metrics of DE$_A$3-guided partial diffusion variants. Candidates with pLDDT > 85, pAE_interaction < 10, and ΔΔG < –30 were selected (red box). Overall, 49.5% of the total designs (49.5%, 3775/7620) passed the filtering criteria. Data points with ΔΔG < –30 are shown in black; those with ΔΔG > –30 are shown in yellow. Among these, final candidates selected for experimental validation (protein expression, purification, and binding analysis) are indicated by red circles. The top 100 candidates indicated in the black box are screened by yeast surface display.

**b.** Yeast surface display screening and FACS-based enrichment of GDF15 binders. In the two-dimensional FACS plot, the x-axis shows GDF15 binding (PE fluorescence) and the y-axis indicates surface displayed binder (PE-Cy7). High-expression and high-binding clones (upper right quadrant, 25.2% of the total population) were isolated for further analysis.

**c.** Structural alignment of DE$_A$3 with DE$_A$3-derived binder variants. RMSD with AF2-predicted structure and AF2 scores (pAE_interaction and pLDDT) for the binder/GDF15 complex are indicated.

**d.** SEC profiles of DE$_A$3-derived binders (DE$_A$3-1, DE$_A$3-2, and DE$_A$3-5).

**e.** SPR analysis of DE$_A$3-1, DE$_A$3-2 and DE$_A$3-5 binding to recombinant GDF15 dimers. Sensorgrams are shown for 5-50 nM analytes (indicated binder).

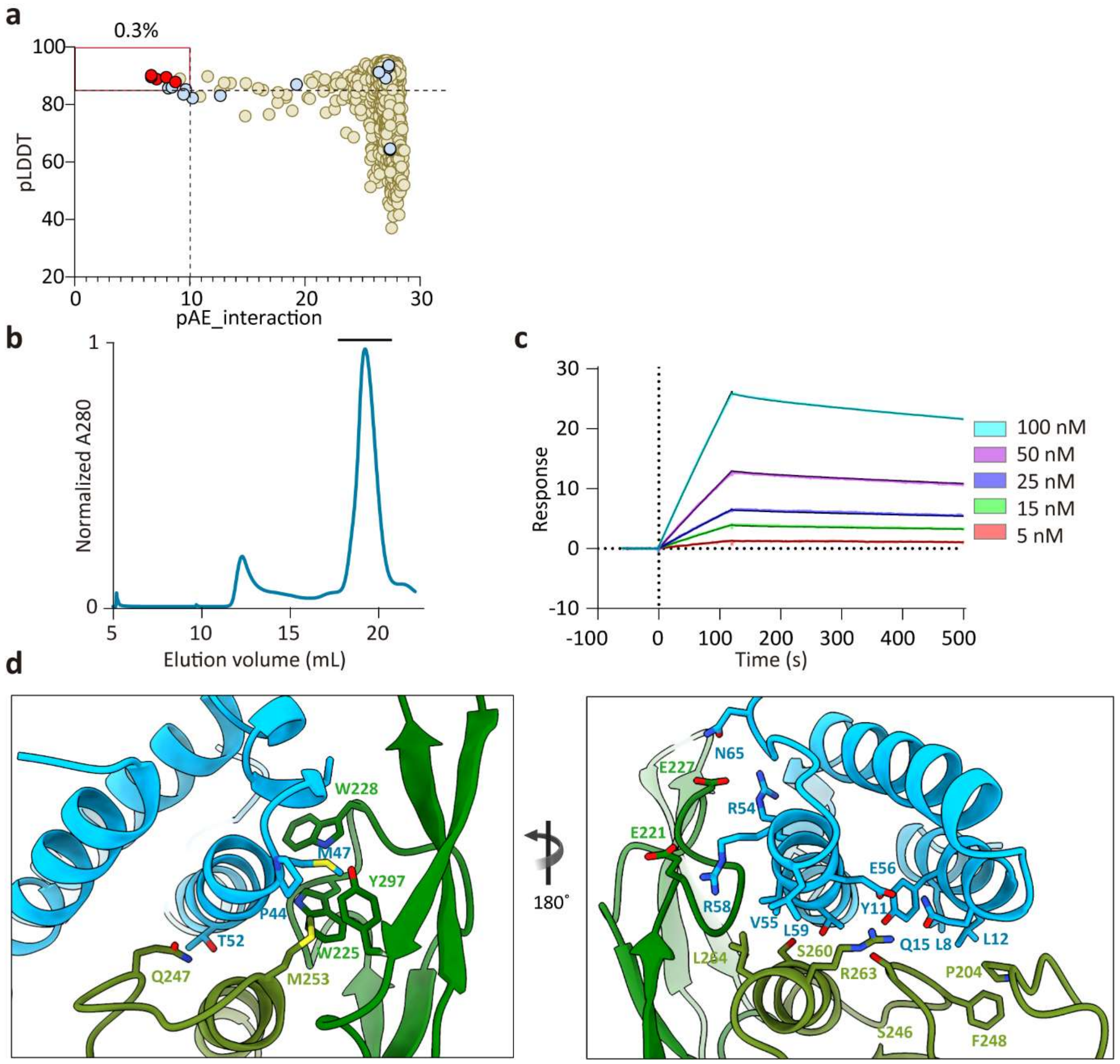


**Figure S4. Structural and biochemical characterization of SSG-designed GDF15 site B binders.**

**a.** Filtering metrics of RGMA-guided partial diffusion variants. Candidates with pLDDT > 85, pAE_interaction < 10, and ΔΔG < –30 were selected (red box). Overall, 0.3% of the total designs (0.3%, 10/3000) passed the filtering criteria. Data points with ΔΔG < –30 are shown in black; those with ΔΔG > –30 are shown in yellow. Among these, final candidates selected for experimental validation (protein expression, purification, and binding analysis) are indicated by red circles.

**b.** SEC profiles of $SSG_B2$ binders.

**c.** SPR analysis of $SSG_B2$ binding to recombinant GDF15 dimers. Sensorgrams are shown for analytes ranging from 5 to 100 nM (indicated binder).

**d.** Binding interfaces between $SSG_B2$ and GDF15 site B. Key interacting residues are shown as sticks and labeled.

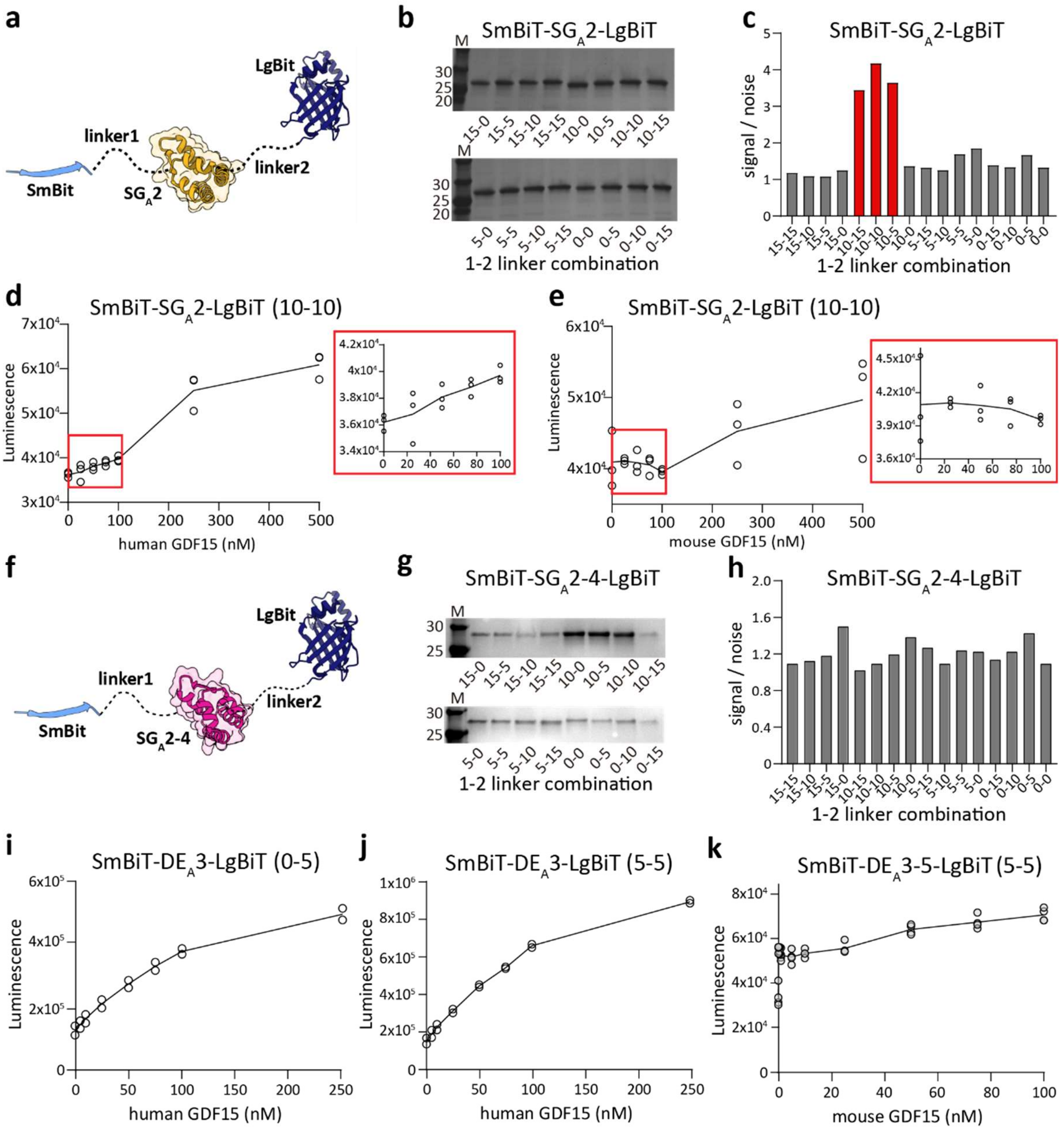


**Figure S5. Linker optimization of BAT biosensor with SG$_A$2 and SG$_A$2-derived GDF15 site A binder.**

**a, f.** Structural model of BAT biosensors constructed with SG$_A$2 **(a)** or SG$_A$2-4 **(f)** binders, flanked by SmBiT at the N-terminus and LgBiT at the C-terminus. The lengths of Linker1 and Linker2 were systematically varied to optimize biosensor performance.

**b, g.** SDS-PAGE analysis of SmBiT–SG$_A$2–LgBiT **(b)** and SmBiT–SG$_A$2-4–LgBiT **(g)** with different linker combinations after *E.coli* expression and affinity purification.

**c, h**. Screening of linker combinations for SmBiT–SG$_A$2–LgBiT **(c)** and SmBiT–SG$_A$2-4–LgBiT **(h)** by luminescence assay. Signal-to-noise ratios (luminescence intensity of each construct divided by that of the control without GDF15) are plotted, with optimal linker combinations for SG$_A$2 highlighted in red.

**d, e.** Luminescent signal of SmBiT–SG$_A$2–LgBiT with 10-10 linkers to human GDF15 **(d)** or mouse GDF15 **(e)**. Luminescence (arbitrary units, AU) is plotted against various GDF15 concentrations (n = 3). The linear detection range is indicated with a red box (0–100 nM).

**i, j.** Luminescent signals of SmBiT–DE$_A$3–LgBiT with 0-5 linkers to human GDF15 **(i)**, with 5-5 linkers to human GDF15 **(j).**

**k.** Luminescent signals of SmBiT–DE$_A$3-5–LgBiT with 5-5 linkers to mouse GDF15.

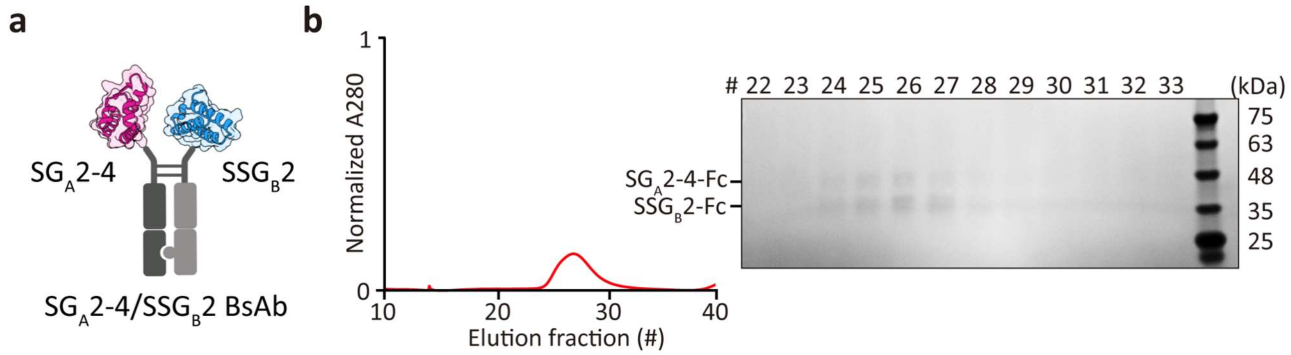


**Figure S6. Linker optimization of BAT biosensor with $SG_A2$ and $SG_A2$-derived GDF15 site A binder.**

**a.** Schematic diagram of Fc-fused $SG_A2$-4/$SSG_B2$ binder (Bispecific $SG_A2$-4/$SSG_B2$-Fc).

**b.** SEC profile of bispecific $SG_A2$-4/$SSG_B2$-Fc (left) and SDS-PAGE analysis of elution fractions (right).
